# Rethinking Syndecan-1 as a Marker of Trauma-Induced Endothelial Injury

**DOI:** 10.64898/2026.10.12.758409

**Authors:** Shibani Pati, Huimin Geng, Alpa Trivedi, Callie Keane, Mark Barry, Praneeti Pathipati, Ken Bertram, Ernest Moore, Martin Schreiber, Joseph Cuschieri, John B Holcomb, Emanuela Zacco, Alison B Nair

## Abstract

Syndecan-1 (SDC1), a transmembrane proteoglycan and component of the glycocalyx, is widely considered the key circulating biomarker of endothelial dysfunction in trauma and critical illness. However, the tissue distribution of SDC1 and the cellular origin of plasma SDC1 in health and disease are incompletely understood. This series of experiments integrates immunostaining, spatial transcriptomics (GeoMx Digital Spatial Profiler (DSP) and Xenium), and single-cell atlases to map SDC1, as well as the other three members of the syndecan family, syndecan-2 (SDC2), syndecan-3 (SDC3), and syndecan-4 (SDC4), across mouse and human tissues, including the lung, gastrointestinal tract, and brain. Across modalities, SDC1 primarily localizes to epithelial compartments, as demonstrated by colocalization with epithelial markers and enrichment in epithelial cell types in the lung and gut. Spatial profiling further identifies SDC1 and SDC4 enrichment in airway epithelial niches, SDC3 in endothelial-rich regions, and SDC2 distributed more broadly across compartments. Mouse and human single-cell atlases confirm epithelial predominance of SDC1. These findings define the cellular and spatial landscape of syndecan expression in healthy tissues and support SDC1 as an epithelial-enriched proteoglycan. These results also challenge the specificity of circulating SDC1 as a biomarker of endotheliopathy and raise the possibility that epithelial injury contributes substantially to circulating SDC1 after trauma and critical illness.

## INTRODUCTION

Trauma triggers a complex, multi-system pathophysiologic response in which the vascular endothelium plays a central role. Composed of a single layer of endothelial cells and associated structures, the vascular endothelium lines blood vessels and regulates critical interactions between the circulating blood and surrounding tissues (1). Following injury, endotheliopathy is driven by local factors, including tissue damage and shear stress, and systemic responses, including cellular dysregulation, immune-mediated injury, and oxidative stress (2–6). The resulting constellation of barrier failure, endothelial dysregulation, coagulopathy, and inflammation characterizes the endotheliopathy of trauma (EOT, 7-9). When severe or persistent, EOT contributes to organ dysfunction, multiple organ failure, and mortality (9–12). Importantly, many of these pathophysiologic mechanisms are shared across other forms of critical illness, suggesting that endothelial dysfunction represents a common pathway contributing to organ injury in diverse disease states.

A key component of endothelial structure and function is the glycocalyx, a dynamic extracellular matrix that regulates vascular homeostasis and is rapidly disrupted in traumatic injury (13–15). The glycocalyx is present on many cell types and tissues including the endothelium, epithelium, peritoneum, and pleura. Following hemorrhagic shock and trauma, glycocalyx degradation is thought to be an inciting event that leads to the cascade of endothelial dysregulation characteristic of EOT (15–27). Although the mechanisms underlying glycocalyx degradation are not fully understood, systemic inflammation, hypoxia, reactive oxygen species, and proteolytic enzymes released by activated leukocytes and injured tissues promote activation of sheddases that cleave glycocalyx constituents from the endothelial surface (16–21,27,28). Loss of the glycocalyx exposes the endothelial surface to mechanical and biochemical stimuli, resulting in endothelial activation and dysfunction (29–32). These changes increase vascular permeability, promote leukocyte and platelet adhesion, amplify inflammatory signaling, and disrupt normal hemostatic regulation (14,15,32–34). The resulting extravasation of plasma into the interstitial space contributes to tissue edema, impaired microvascular perfusion, and organ dysfunction, while activation of inflammatory, coagulation, and immune pathways further perpetuates endothelial injury (6,24,35–39).

Despite the central role of endotheliopathy in trauma, tools to directly assess specific endothelial injury remain limited. Consequently, circulating biomarkers of glycocalyx disruption have emerged as attractive surrogates given the glycocalyx’s early involvement in endothelial dysfunction. Among these, syndecan-1 (SDC1), a transmembrane heparan sulfate proteoglycan, was the first syndecan to be easily measured, and has thus become one of the most extensively studied circulating biomarkers of EOT (40–47). SDC1 belongs to a family of four transmembrane proteoglycans, which also includes syndecan-2 (SDC2), syndecan-3 (SDC3), and syndencan-4 (SDC4), that contribute to glycocalyx structure and regulate cell signaling, adhesion, and interactions with the extracellular matrix (14,15,39). Circulating SDC1 concentrations increase following traumatic injury and have been associated with polytrauma, traumatic brain injury, blunt trauma, and hemorrhagic shock (2,10,38,48–55). In injured patients, elevated SDC1 levels also correlate with injury severity and outcomes, including organ dysfunction and mortality (2,22,26,41–43,50,56–69). Beyond trauma, increased circulating SDC1 has been associated with disease severity and adverse outcomes in sepsis, burns, acute respiratory failure, and other forms of critical illness (37,44,70–89).

A fundamental limitation of SDC1 as a biomarker is its uncertain cellular origin. Although circulating SDC1 is widely interpreted as a marker of endothelial injury, it is also expressed in multiple non-endothelial tissues, raising important questions about its specificity in the circulation (90–92). Consequently, it remains unclear whether elevated plasma SDC1 truly reflects EOT or injury to other cellular compartments. This distinction is important because SDC1 is expressed by epithelial cells, which form barrier surfaces across multiple organs that are themselves vulnerable to injury during trauma and critical illness. Thus, interpreting circulating SDC1 as evidence of endothelial glycocalyx injury assumes an endothelial origin that has not been definitively established. Accurate phenotyping of EOT and, more broadly, endothelial dysfunction is critical, as it plays a role in common sequelae of severe illness including shock, capillary leak, coagulopathy, dysregulated inflammation, and multiple organ dysfunction (23,93). Furthermore, the development of robust, clinically accessible biomarkers that accurately define the underlying cellular pathology is essential to improve risk stratification, identify therapeutic targets, and advance precision therapies for trauma and critical illness (94).

Recent advances in spatial and single-cell transcriptomics enable high-resolution systematic mapping of gene expression while preserving cellular and anatomic architecture. These approaches provide an unprecedented opportunity to define the tissue- and cell type-specific distribution of SDC1 and related syndecans (SDC2-4) across organs and species. This series of experiments integrates immunostaining, spatial transcriptomics (GeoMx Digital Spatial Profiling (DSP) and Xenium), and large-scale single-cell atlases to characterize the cellular and spatial landscape of syndecan expression in mouse and human tissues. By resolving syndecan localization across epithelial, endothelial, mesenchymal, and immune compartments, we aim to clarify the biological origin of circulating SDC1 and refine its interpretation as an endotheliopathy biomarker. We hypothesized that SDC1 would not be restricted to the endothelial compartment and that defining its cellular distribution would provide a more biologically informed framework for interpreting circulating SDC1 in trauma and critical illness.

## RESULTS

### SDC1 and epithelial cell adhesion molecule (EpCAM) demonstrated colocalization in the mouse lung, small intestines, and brain

Colocalization of SDC1 was evaluated in the mouse lung, small intestine, and brain by immunostaining for the endothelial-specific vascular endothelial cadherin (VE-cadherin) and the epithelial-specific EpCAM (**Figure 1**). In all sections, SDC1 and VE-cadherin showed minimal overlap, with distinct localization of each in the merged images (**Figure 1A**). In contrast, SDC1 and EpCAM exhibited colocalization, as shown in yellow in the merged images. SDC1 was localized to small airways in the lung, microvilli of the small intestine, and the brain’s choroid plexus, all structures where epithelial cells are present (**Figure 1B**).

**Figure 1:**
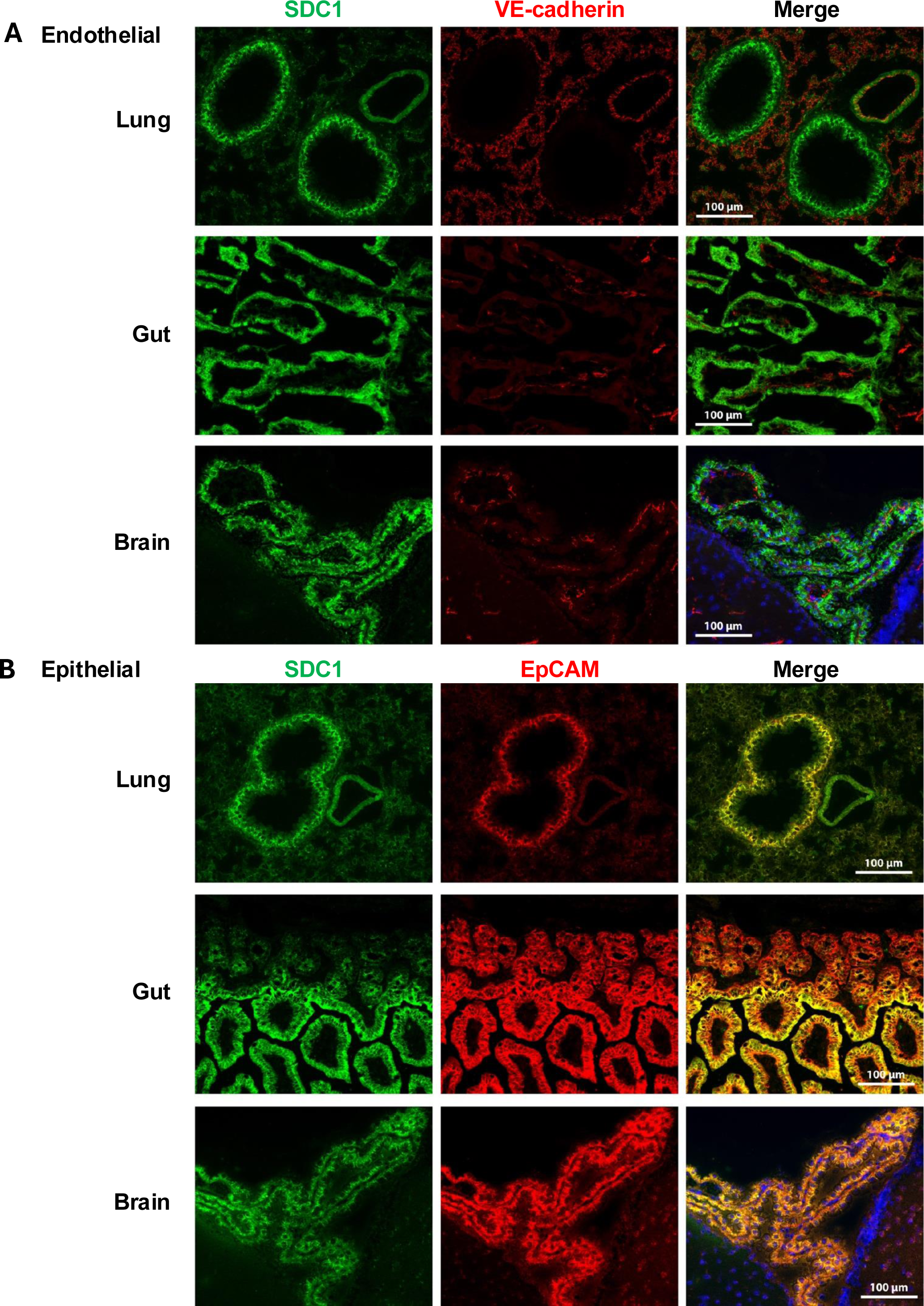
Immunoexpression of Syndecan-1 (SDC1) protein primarily on epithelial cells across multiple tissues. (**A**) Sections of the lung, brain, and gut were stained with antibodies against SDC1 (green) and vascular endothelial-cadherin (VE-cadherin, red), and images were merged. There was no overlapping staining between VE-cadherin and SDC1. 4′,6-diamidino-2-phenylindole (DAPI, blue) was added to the brain merged images. (**B**) Sections of the lung, brain, and gut were stained with antibodies against SDC1 (green) and epithelial cell adhesion marker (EpCAM, red), and images were merged. Overlapping areas in the merged images appear yellow. DAPI staining was added to the merged brain images.

### GeoMx DSP identified epithelial-enriched SDC1 expression in the mouse lung

Given the qualitative overlap of SDC1 with the epithelial marker EpCAM, spatial transcriptomics by GeoMx DSP was performed to quantify SDC1 presence in the epithelial and endothelial compartments of the mouse lung. A total of 69 regions of interest (ROIs) were selected for three lung niches, specifically 36 from the airway niche, 20 from the blood vessel niche, and 13 from the alveoli niche (**Figure 2A-2C, Supplemental Figure 1**). Uniform manifold approximation and projection (UMAP) distribution revealed separation of the three niches (**Figure 2D)**. Abundance scores indicated a predominance of epithelial cells in the airway niche, smooth muscle cells in the blood vessel niche, and endothelial cells in the alveoli niche (**Figure 2E**). Spatial deconvolution based on single-cell signature abundance scores for each ROI across lung compartments further demonstrated epithelial, smooth muscle (SM), and endothelial dominance of the airway, blood vessel, and alveoli niches, respectively (**Figure 2F**).

**Figure 2:**
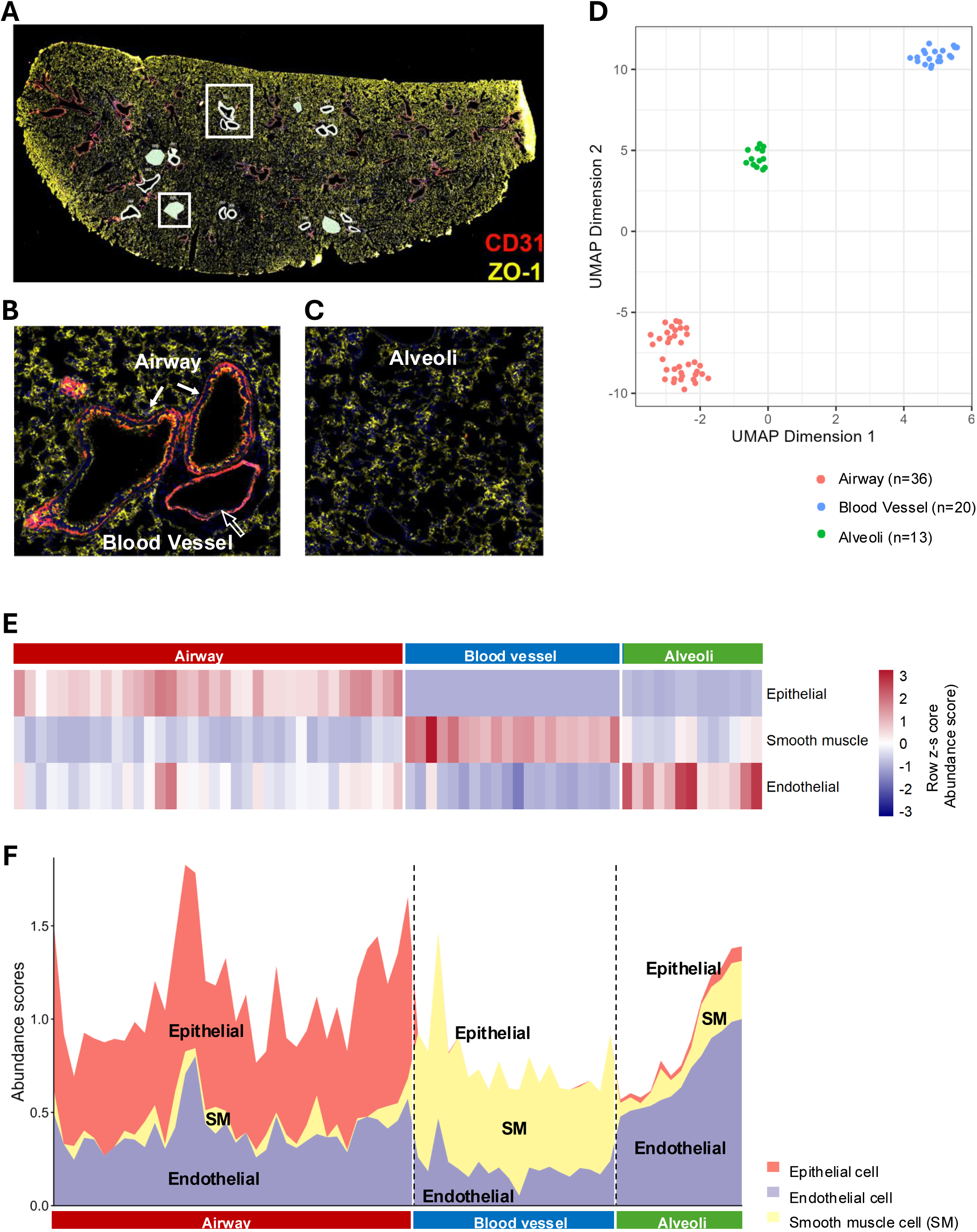
Spatial transcriptomic profiling of lung niches using GeoMx Digital Spatial Profiling (DSP). **(A)** Low-magnification image of the lung section stained with the morphology markers CD31/platelet endothelial cell adhesion molecule-1 (PECAM1, red, endothelial marker), zonula occludens-1 (ZO-1, yellow, tight junction marker), and DAPI (blue). These markers highlight endothelial, epithelial, and vascular structures to guide precise region of interest (ROI) placement. The selected ROIs are outlined in white. White boxes indicate two representative ROIs displayed in panels (B) and (C). (**B-C**) Digitally zoomed image of the ROI is shown to represent the airways, blood vessels **(B)**, and alveoli **(C)**. **(D)** Uniform manifold approximation and projection (UMAP) embedding of the selected ROIs demonstrates distinct clustering according to lung compartment: airway, blood vessel, and alveoli. **(E)** Heatmap shows scaled abundance scores of major cell types (epithelial cells, smooth muscle cells, and endothelial cells) across the three lung tissue compartments (airway, blood vessel, alveoli). Each column represents an individual ROI, ordered by airway, blood vessel, and alveoli, and each row represents a cell-type signature. Color bar denotes scaled abundance scores. **(F)** Spatial deconvolution of cell types based on single-cell signature abundance scores for each ROI across lung compartments. Stacked abundance plots display estimated proportions of the major cell types, epithelial cells, endothelial cells, and smooth muscle (SM), for each ROI. Samples are grouped by lung niches (airway, blood vessel, alveoli). Vertical dashed lines mark compartment boundaries.

After establishing ROIs, expression patterns of SDC1-4 were evaluated in the epithelial and endothelial compartments as well as across the airway, blood vessel, and alveoli niches. Both SDC1 and SDC4 were more highly expressed in epithelial regions, whereas SDC3 demonstrated more endothelial expression, and SDC2 was more evenly expressed in both compartments (**Figure 3A-B**). Spearman correlation analysis yielded a strong, statistically significant correlation of SDC1 gene expression in epithelial cells (**Figure 3B**). SDC4 and SDC3 showed weaker but still statistically significant correlations with expression in epithelial and endothelial cells, respectively. SDC2 expression was not statistically correlated with either cell type (**Figure 3B**).

**Figure 3:**
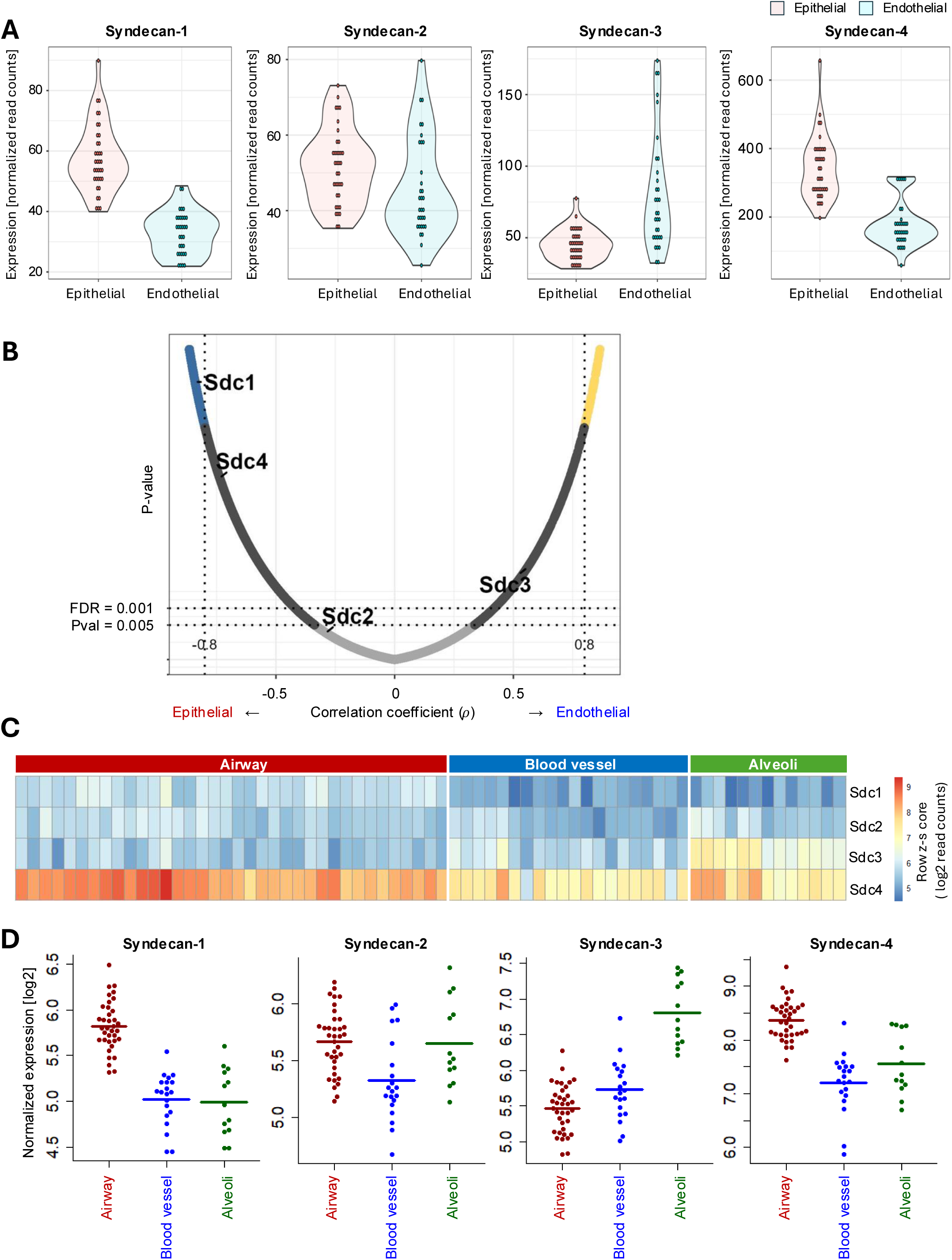
Expression patterns of syndecan genes in epithelial versus endothelial lung compartments. **(A)** Violin plots show normalized expression levels for each syndecan family member in epithelial-dominant (red) and endothelial-dominant (blue) lung regions. Each point represents an individual ROI, and the shape of the violin indicates the kernel density of the data. Each ROI was classified as epithelial-dominant if epithelial cells comprised more than 60% of the composition, endothelial-dominant if endothelial cells exceeded 60%, or excluded from analysis otherwise. SDC1 and syndecan-4 (SDC4) are more highly expressed in epithelial regions, while syndecan-3 (SDC3) shows elevated expression in endothelial compartments. **(B)** Spearman correlation analysis was performed between SDC1-4 gene expression and cell types (epithelial or endothelial), reporting the correlation coefficient (ρ), and P-values. In the plot, each point is a gene. The x-axis shows the Spearman correlation coefficient (ρ), with negative values indicating association with epithelial-dominant ROIs and positive values indicating association with endothelial-dominant ROIs. The y-axis represents statistical significance. SDC1 and SDC4 were associated with epithelial-dominant ROIs, whereas SDC3 was associated with endothelial-dominant ROIs; Syndecan-2 (SDC2) showed a weaker, intermediate association. Vertical dashed lines indicate correlation-coefficient thresholds of ρ = −0.8 and ρ = 0.8. Horizontal dashed lines indicate the specified significance thresholds of P = 0.005 and false discovery rate (FDR) = 0.001. **(C)** Heatmap of SDC1, SDC2, SDC3, and SDC4 gene expression in each ROI across lung niches, including airway, blood vessel, and alveoli regions. Each column represents an individual ROI, and each row corresponds to a gene. The color bar reflects normalized expression levels. **(D)** Scatter plots summarize expression differences of SDC1, SDC2, SDC3, and SDC4 across airway, blood vessel, and alveoli ROIs. Each dot represents an ROI, and the horizontal bars indicate group means. SDC1 and SDC4 are more abundant in the airway, whereas SDC3 is most enriched in the alveoli. SDC2 exhibits a more uniform distribution.

Gene expression of SDC1-4 across the airway, blood vessel, and alveoli niches further elucidates SDC location and distribution (**Figure 3C-3D**). Overall, SDC4 was the most highly expressed across niches, and SDC1 was the lowest. Both SDC1 and SDC4 had more prominent expression and abundance in the airway niche compared to the other niches. SDC3 was more enriched in the alveoli niche, and SDC2 exhibited a more uniform distribution. Taken together, based on GeoMx DSP spatial transcriptomics, SDC1 was most abundant in the mouse airway, which contained a dominance of epithelial cells.

### Xenium spatial profiling showed SDC1 is primarily epithelial in the mouse embryo

Xenium spatial transcriptomics was used to evaluate SDC1 more broadly in a sagittal mouse embryo. SDC1 expression displayed the highest expression density seen in the skin, oropharynx, and whiskers (**Figure 4A-B**). At higher magnification, SDC1 transcripts were present in both the epithelial and stromal compartments indicated by overlay with EpCAM and keratin 8 (KRT8) transcripts **(Figure 4C-D)**. SDC1 was distinct from the endothelial compartment, with transcripts differentially located from the endothelial von Willebrand Factor (VWF, **Figure 4E-F**).

**Figure 4:**
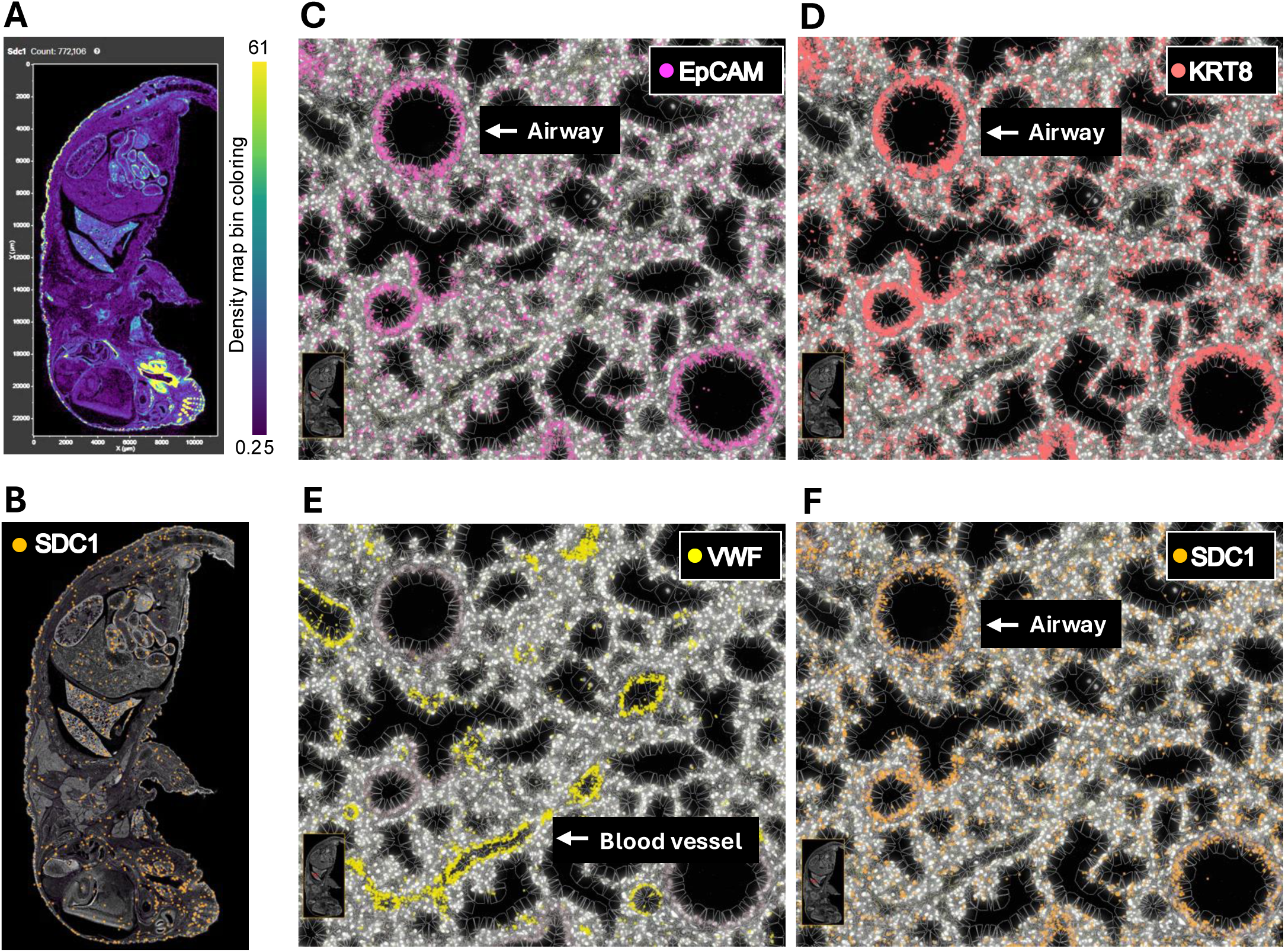
Spatial expression of SDC1 in a sagittal mouse embryo section in the Xenium Mouse Multi-Tissue Panel. (A-B) Low-magnification overview of SDC1 spatial expression across a full sagittal section of a developing mouse embryo (whole body, formalin-fixed, paraffin-embedded). The upper panel displays a density map of SDC1 expression, with the color scale ranging from low (purple) to high (yellow, **A**). In the lower panel, each dot represents a detected SDC1 transcript mapped to a spatial location within the whole-body section. The grayscale background image shows tissue morphology (**B**). **(C-F)** High-magnification view of the spatial transcriptomic map in a region of the embryonic mouse lung acquired using the 10x Genomics Xenium platform. Transcripts for EpCAM (pink, **C**) and Keratin 8 (KRT8, red, **D**) indicate epithelial cells; von Willebrand Factor (VWF, yellow, **E**) highlights endothelial cells; SDC1 (orange, **F**) colocalizes with epithelial and stromal compartments. Cell boundaries are outlined in white based on segmentation data. The background shows tissue morphology in grayscale.

### Single-cell LungMAP confirmed predominant epithelial expression of SDC1 in the mouse lung

To further characterize SDC1 expression in the lung, SDC1 expression was mapped onto the mouse lung cellular profile. Using consolidated single-cell transcriptomic data from a cohort of mouse lung samples, a cellular reference MAP was constructed (**Figure 5A, Supplemental Figure 2**). SDC1 expression by cell type was primarily present in epithelial subpopulations, including AT1, AT2, ciliated, and secretory cell types (**Figure 5B-5C**). SDC4 also demonstrated a predominantly epithelial overlay, with expression in epithelial cell types similar to SDC1. SDC2 was more highly expressed in mesenchymal cell types. SDC3, which had lower overall expression than the other SDCs, was expressed most in endothelial cell types (**Figure 5C**).

**Figure 5:**
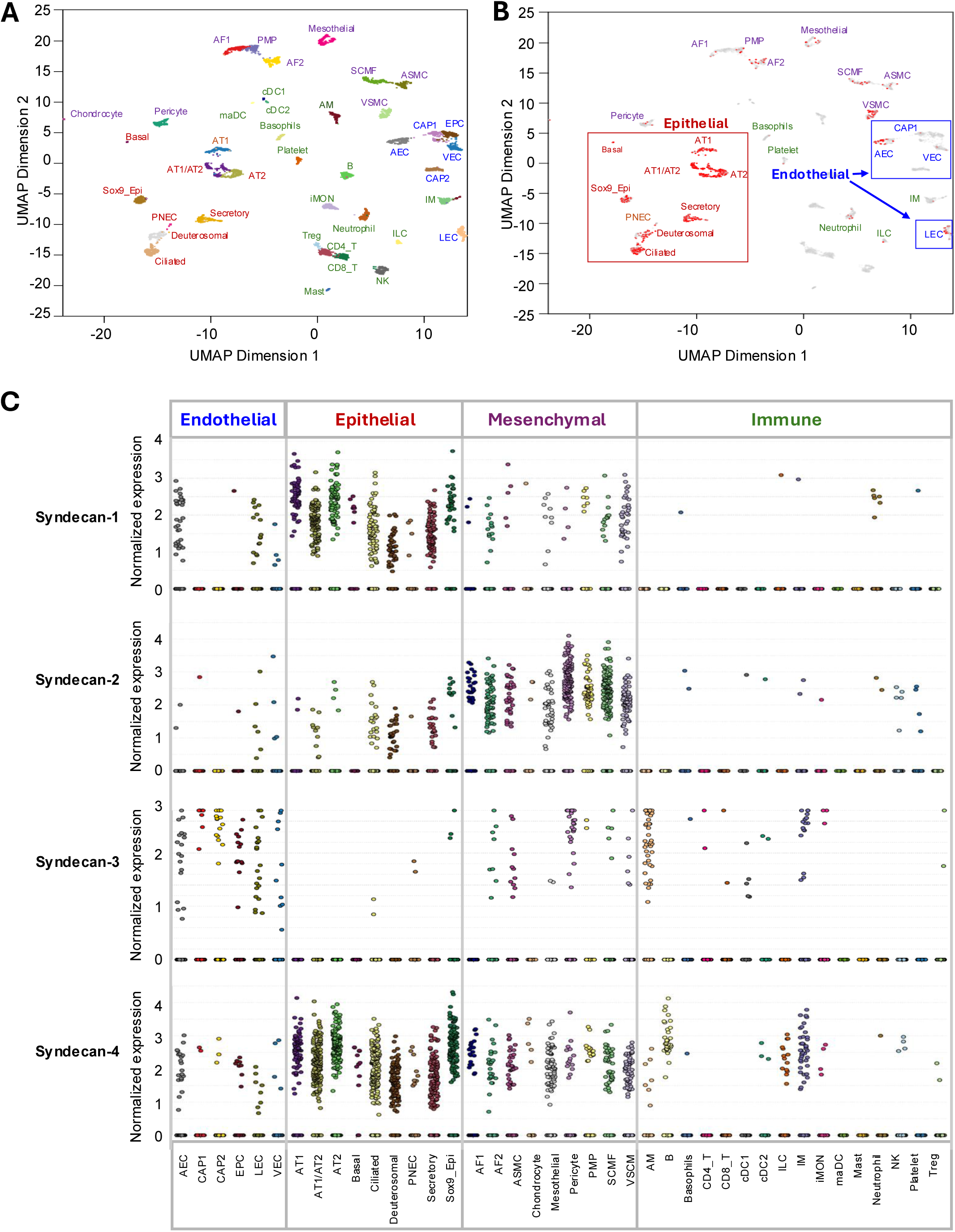
Syndecan expression in mouse lung cells from the Lung MAP. **(A)** UMAP visualization of 40 annotated cell types (clusters) from mouse lungs from the Lung MAP, Lung Gene Expression Analysis (LGEA) web portal (see **Supplemental Figure 2** for details). Each cluster is colored by its predicted cell identity. **(B)** Visualization of SDC1 gene expression mapped onto the UMAP shown in panel A. Colors represent the SDC1 expression, with red indicating detectable SDC1 expression, primarily within epithelial subpopulations, and gray indicating u ndetectable SDC1 expression. **(C)** Expression profiles of SDC1, SDC2, SDC3, and SDC4 across mouse lung cell types, summarized by four major categories: epithelial, endothelial, mesenchymal, and immune cells.

### SDC1 expression was preferentially epithelial in the human lung

To characterize SDC1 in human tissues, references for the human lung, gastrointestinal system, and brain were interrogated for its expression. Using consolidated single-cell transcriptomic data from a cohort of human lung samples, a cellular reference MAP was constructed (**Figure 6A, Supplemental Figure 3**). As in the mouse lung, SDC1 expression by cell type demonstrated a predominance in epithelial subpopulations. Expression in plasma cells was also noted (**Figure 6A-6B**). As in the mouse lung, SDC4 expression was strongest in epithelial cells, SDC2 expression was strongest in mesenchymal cells, and SDC3 was present across cell populations, but less strongly than the other SDCs. While the overall distribution across epithelial, endothelial, mesenchymal, and immune cells was similar between the mouse and human lung, there was notable variation in gene expression by cell type within each category (**Figure 6C**).

**Figure 6:**
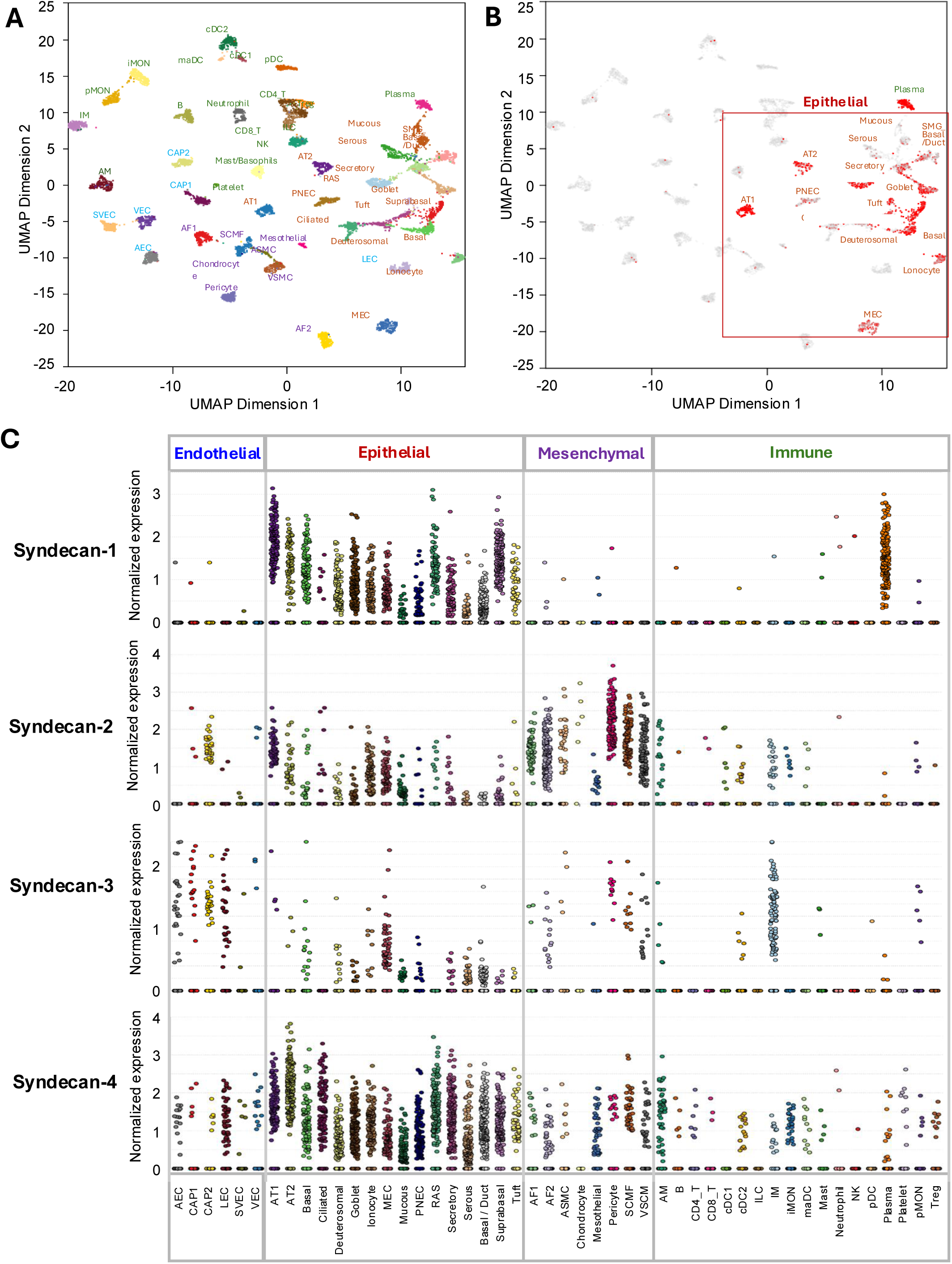
SDC expression in normal human lung cells from the Lung MAP. **(A)** UMAP visualization of 48 annotated cell types (clusters) from normal human lungs, based on the Lung MAP, Human Lung CellRef Seed (see **Supplemental Figure 3** for details). Each cluster is colored by its predicted cell identity. **(B)** Visualization of SDC1 gene expression mapped onto the UMAP shown in panel A. Colors represent the SDC1 expression, with red indicating detectable SDC1 expression, primarily within epithelial subpopulations, and gray indicating undetectable SDC1 expression. **(C)** Expression profiles of SDC1, SDC2, SDC3, and SDC4 across human lung cell types, summarized by four major categories: epithelial, endothelial, mesenchymal, and immune cells.

### Genotype-tissue expression project (GTEx) bulk and single-cell analyses showed epithelial-enriched SDC1 expression across human tissues

Analysis of GTEx bulk RNA sequencing data revealed that SDC1 expression is enriched in epithelial-associated tissues, with the highest transcript levels observed in stratified and mucosal epithelia and comparatively low expression in vascular-associated tissues (**Supplemental Figure 4A**). Consistent with this pattern, single-cell RNA sequencing data demonstrated that SDC1 expression is concentrated in epithelial cell populations across tissues, with minimal signal in endothelial compartments (**Supplemental Figure 4B**). Together, these results support a predominant epithelial localization of SDC1 across human tissues.

In contrast, SDC2–4 exhibited distinct and complementary expression patterns. Bulk RNA-seq analysis showed broader tissue distribution of SDC2 and consistent epithelial enrichment of SDC4 (**Supplemental Figure 5A-C**). Single-cell RNA sequencing further resolved lineage-specific partitioning, with SDC2 enriched in mesenchymal compartments, SDC3 associated with endothelial populations, and SDC4 enriched in epithelial cells (**Supplemental Figure 6A-C**). Together, these data highlight lineage-specific syndecan partitioning and support a conserved epithelial bias for SDC1 expression.

### SDC1 expression was primarily epithelial in the human gastrointestinal tract

SDC1 expression patterns in the human gut also revealed selective enrichment for epithelial and plasma cells (**Supplemental Figures 7,8**). As in the lung, the strongest expression was observed in epithelial cells and B plasma cells, while endothelial cells showed minimal SDC1 expression (**Supplemental Figure 8A-B**). Overlay of components of the gastrointestinal tract demonstrates differential presence of SDC1 and suggests much of its expression may be localized to the esophagus and other upper gastrointestinal sources (**Supplemental Figure 8C**). The expression pattern of SDC2 was primarily mesenchymal, SDC4 was primarily epithelial, and SDC3 was distributed across cell types (**Supplemental Figure 9A-F**).

### SDC1 expression is minimal in human brain

SDC1 expression in the human brain was negligible. Specifically, SDC1 expression across neuronal clusters, including neuronal samples across a broad range of anatomical structures, was minimal (**Supplemental Figure 10, 11A-C**). Non-neuronal cells, including fibroblasts, endothelial cells, astrocytes, oligodendrocytes, and pericytes, also demonstrated a uniformly low to absent expression in non-neuronal lineages (**Supplemental Figure 11D-G**). SDC2 and SDC4 did not demonstrate detectable enrichment in either neuronal or non-neuronal cell types. SDC3, however, showed low but detectable expression in subclusters of neuronal and non-neuronal cells (**Supplemental Figure 12A-H**).

## DISCUSSION

Through integrated spatial transcriptomic, single-cell transcriptomic, and immunohistochemical analyses, this study comprehensively defined the tissue- and cell type-specific distribution of SDC1-4 across the lung, gastrointestinal tract, and brain in both mice and humans. Across complementary experimental platforms, SDC1 consistently localized to epithelial compartments rather than endothelial cells. In the mouse lung, this was shown through colocalization with epithelial markers, enrichment in epithelial regions by spatial transcriptomics, and preferential expression in epithelial cell populations by single-cell analyses. Human lung and gastrointestinal tissues demonstrated a remarkably similar pattern, confirming that the epithelial predominance of SDC1 is conserved across species. In both organs, SDC1 expression was concentrated within epithelial cell types, with additional expression identified in plasma cells. Together, these findings demonstrate that SDC1 is an epithelial-enriched proteoglycan and is not specific to the endothelial compartment.

The remaining syndecans demonstrated distinct but overlapping expression patterns. Like SDC1, SDC4 was predominantly expressed in epithelial compartments, suggesting a potential shared role in epithelial barrier biology. In contrast, SDC2 and SDC3 exhibited broader expression across multiple cellular compartments, including endothelial, mesenchymal, and immune populations. Notably, SDC3 showed relative enrichment in endothelial-rich regions of the lung, suggesting that individual syndecans may differentially reflect specific cellular compartments. Although additional functional studies are needed, these observations suggest that different syndecans may provide complementary information regarding injury to distinct tissue compartments rather than serving as interchangeable biomarkers of endothelial dysfunction (44,45,95).

These findings have important implications for interpreting circulating SDC1 following traumatic injury and critical illness. While plasma SDC1 has traditionally been viewed as a marker of endothelial glycocalyx degradation and endothelial injury, our data demonstrating epithelial predominance suggest this interpretation may be incorrect and that circulating SDC1 should not be considered endothelial-specific. Elevations in circulating SDC1 may arise from epithelial injury, endothelial injury, or a combination of injured cellular compartments, with their relative contribution dependent on the underlying disease process, the organs involved, and the timing of disease progression. This interpretation provides a potential explanation for the broad association of circulating SDC1 with diverse forms of trauma and critical illness, including hemorrhagic shock, sepsis, acute respiratory distress syndrome, and burns, all of which involve substantial epithelial barrier disruption in addition to endothelial dysfunction. Further, the intestinal epithelium, which may be uniquely sensitive to hypoxemia and resulting organ dysfunction in critical illness and injury, may emerge as an important contributor to epithelial-associated plasma SDC1 in this context (40,46,96).

Another important observation was the minimal expression of all syndecans in normal brain tissue. These findings suggest that circulating syndecans likely do not originate directly from brain parenchyma and may therefore have limited utility as biomarkers of brain tissue injury. Instead, elevated plasma SDC1 following traumatic brain injury could reflect systemic epithelial or endothelial injury accompanying trauma, although injury-induced changes in cerebral SDC1 expression or shedding cannot be excluded (51,54,89).

This study leverages recent advances in spatial and single-cell transcriptomics to characterize syndecan expression while preserving both cellular identity and tissue context. Recognizing the strengths and limitations of each approach, this work integrated complementary methods to validate findings across independent experimental platforms. The consistency observed across these modalities provides a robust framework for defining the cellular and spatial distribution of syndecans in healthy tissues. Beyond refining our understanding of syndecan biology, this work establishes an important reference atlas for future studies of traumatic injury and critical illness. By providing a foundation for interpreting tissue- and cell type-specific biomarker expression, these findings may facilitate the development of more precise biomarker approaches that distinguish distinct injury phenotypes and ultimately guide targeted therapeutic strategies.

Collectively, these findings challenge the prevailing interpretation of SDC1 as a specific biomarker of endotheliopathy and instead support a more nuanced model in which circulating SDC1 reflects injury across multiple tissue compartments, particularly the epithelium (91,92,97). Rather than diminishing the clinical value of SDC1, these results refine its biological interpretation and underscore the importance of understanding tissue-specific biomarker origins. More broadly, these findings highlight the need to move beyond single biomarkers toward integrated biomarker approaches that distinguish endothelial, epithelial, immune, and mesenchymal injury, providing a more precise framework for phenotyping trauma and critical illness and identifying targeted therapeutic strategies.

While the importance of endotheliopathy is recognized, epithelial injury has received comparatively little attention despite its critical role in maintaining barrier integrity and organ function (91,92,100). Disruption of epithelial barriers contributes to edema, inflammation, susceptibility to infection, and organ dysfunction across multiple organ systems (100–105) and may represent an underappreciated contributor to adverse outcomes following injury. Importantly, these findings highlight a major gap in the field, namely that there are currently no validated biomarkers that specifically quantify endothelial injury, distinguish clinically meaningful endotypes of trauma, or guide precision-based therapeutic interventions. Likewise, reliable biomarkers of epithelial injury remain lacking. Addressing these gaps will require the development of biomarkers that more accurately reflect the specific underlying biology of injury.

Importantly, one can ask how proteins like Syndecan-1, which are largely expressed in epithelial cells, can reach the systemic circulation. Cells within peripheral tissues, including epithelial, stromal, and resident or infiltrating immune cells, continuously secrete a diverse repertoire of soluble proteins, including cytokines, chemokines, growth factors, enzymes, extracellular-matrix–associated proteins, and other signaling mediators. Once released, these factors enter the interstitial fluid, where their distribution depends on molecular size, extracellular-matrix interactions, vascular permeability, and interstitial fluid flow. Smaller soluble molecules may easily access the blood microvasculature, whereas larger proteins and macromolecular complexes can be transported through the lymphatic circulation. Blind-ended lymphatic capillaries efficiently take up interstitial fluid and tissue-derived proteins and transport this material through collecting lymphatics to regional lymph nodes. In this way, lymph provides a molecular “snapshot” of the secretory and inflammatory state of the tissue it drains (99). After passage through lymph nodes, lymph and its soluble protein cargo enter the venous circulation via the thoracic and right lymphatic ducts, providing a direct mechanism by which locally produced proteins from epithelial cells can become systemically distributed. Tissue injury, inflammation, and various resuscitation interventions may further amplify this process by increasing vascular permeability and altering lymphatic flow, thereby enhancing the movement of tissue-derived mediators into the systemic circulation (99).

Importantly, this study has several limitations. Both spatial and single-cell transcriptomic approaches are inherently dependent on the tissues and samples analyzed. As a result, these approaches may not fully capture biological variability across individuals. Specifically, GeoMx DSP interrogates regions of interest containing multiple cells rather than individual cells, resulting in aggregated gene expression signals that are influenced by tissue composition and ROI selection. Furthermore, cell-type deconvolution relies on reference transcriptomes and assumes stable transcriptional profiles, which may not fully reflect biological heterogeneity. Similarly, single-cell transcriptomic analyses are limited by the number of cells and individuals included in the reference datasets. Importantly, syndecan expression likely varies across individuals, ages, and disease states, and the extent of this variation remains unknown. Although concordance across complementary methodologies strengthens our conclusions, these observations should be validated in additional cohorts and tissues.

An additional limitation is that this study characterizes syndecan expression in healthy tissues and therefore does not directly establish the cellular source of circulating SDC1 after traumatic injury. Moreover, transcript abundance and tissue localization do not necessarily predict protein abundance or the magnitude of protein shedding into the circulation following injury. Syndecan shedding is likely a dynamic process shaped by the host, the injury, the host’s response, and treatment variability (65,73,87,88,98), and the relative contributions of epithelial, endothelial, immune, and other cellular compartments remain unknown. Defining the mechanisms and cellular origins of SDC1 shedding after injury is an important area for future investigation and will be critical for refining the interpretation of circulating SDC1 as a biomarker of trauma and critical illness.

In conclusion, trauma-related endotheliopathy remains a critical driver of the pathophysiology that follows injury. Earlier studies often interpreted circulating SDC1 primarily as a marker of endothelial glycocalyx degradation and, consequently, as a surrogate for endothelial injury. However, advances in single-cell RNA sequencing and spatial transcriptomics, as utilized in this paper, have substantially refined our understanding of the cellular distribution of SDC1. This shift reflects the normal evolution of scientific thought as techniques to interrogate molecular and cellular function become more precise. These approaches demonstrate that SDC1 is expressed predominantly by epithelial cell populations in many tissues, rather than being an endothelial-specific protein. Thus, elevations in circulating SDC1 should not necessarily be interpreted as evidence of endothelial injury alone but may also reflect epithelial glycocalyx disruption and tissue injury. This distinction is particularly important in complex systemic conditions such as trauma, critical illness, and other inflammatory conditions, where both epithelial and endothelial compartments may contribute to circulating SDC1.

## METHODS

### Animal protocol

Animal studies were performed with approval of the Institutional Animal Care and Use Committee at the University of California, San Francisco (UCSF). The experiments complied with the National Institutes of Health guidelines on the use of laboratory animals. All animals were housed in a room with ad libitum access to food and water, controlled temperature, and a 12:12-hour light-dark cycle.

### Tissue processing, immunostaining, and imaging

Organs from mice were excised, flash-frozen in 100% isopentane, and stored at −80°C, as described previously (106). Flash-frozen brains, lungs, and intestines were sectioned at 10 µm on a Leica CM 1950 Cryostat (Leica Biosystems). Fresh-frozen tissue sections were fixed in 95% ethanol and acetone, then stained with antibodies against SDC1 (Abcam), VE-cadherin (RCD Systems), and EpCAM (Abcam). Fluorescently tagged secondary antibodies were added for detection and imaging. The stained sections were imaged with a Nikon Eclipse 80i microscope (Nikon Instruments Inc) using a SPOT RT-sCMOS camera (SPOT Imaging) at 20x magnification.

### GeoMx DSP image processing and ROI selection

Mouse lung was embedded in optimal cutting temperature (OCT) compound, sectioned at 8 µm thickness, and collected on SuperFrost Plus slides (Thermo Fisher Scientific). Spatial transcriptomic profiling was performed using the GeoMx DSP (Bruker Spatial Biology) according to the manufacturer’s protocols and standard instrument parameters. Tissue sections were hybridized with the GeoMx Whole Transcriptome Atlas (WTA) probe panel and stained with fluorescent morphology markers to guide compartment-specific region selection. The morphology panels included STYO13 for nuclear staining and antibodies against CD31 and ZO-1.

High-resolution fluorescence images were acquired and reviewed within the GeoMx DSP system. ROIs representing airway, alveoli, and blood vessel compartments were selected based on histologic architecture and immunofluorescent marker expression. After ROI selection, ultraviolet illumination was directed to each ROI using the instrument’s programmable digital micromirror device, releasing photocleavable oligonucleotide tags from the hybridized probes. The released oligonucleotides were collected into individual wells of a microtiter plate for downstream library preparation. Tissue processing, staining, ROI collection, and library preparation were performed by the UCSF LCA-Genome core. An overview of the GeoMx DSP workflow is provided in **Supplemental Figure 1**.

### Library preparation, sequencing, and data preprocessing

Collected oligonucleotide tags were amplified by PCR to generate indexed sequencing libraries according to the GeoMx next-generation sequencing (NGS) library preparation protocol. Libraries were pooled and sequenced on the Illumina NextSeq platform. FASTQ; files were processed using the NanoString GeoMx NGS Pipeline. Read trimming, unique molecular identifier collapsing, and probe-to-target mapping were performed according to manufacturer recommendations. ROIs were retained if they met the following quality control criteria: sequencing saturation greater than 50%, more than 1,000 raw reads, alignment rate greater than 80%, geometric mean of negative probe count greater than 10, more than 50 nuclei, median target-probe signal exceeding the mean negative-control probe signal by at least 2 standard deviations, and fewer than 20% of probes flagged as low-count or outlier. ROIs that failed to meet these criteria were excluded from downstream analyses. After quality control, count matrices were normalized using third-quartile normalization in the NanoString GeoMx analysis software. The normalized expression matrices were exported for downstream analysis in R.

### Cell type deconvolution and data visualization

Cellular composition within each ROI was estimated using the SpatialDecon R package (NanoString Technologies), a log-normal regression–based deconvolution framework optimized for GeoMx DSP data. SpatialDecon models ROI-level gene expression as a linear combination of cell type-specific reference profiles. Reference expression matrices were obtained from the NanoString “Human_Cell_Landscape” dataset. Heatmaps and related visualizations were generated using the R packages ggplot2 and pheatmap. Normalized expression values for SDC1, SDC2, SDC3, and SDC4 were extracted for each ROI. Scatter plots were generated to compare SDC1-4 expression levels across anatomical compartments, including airway, alveoli, and blood vessel regions. Abundance scores derived from spatial deconvolution were used to estimate the relative proportions of epithelial and endothelial cells within each ROI. Based on these estimates, ROIs were classified into two categories: predominantly epithelial and predominantly endothelial, according to the dominant cell-type signature. Violin plots were generated to compare the expression levels of SDC1, SDC2, SDC3, and SDC4 between predominantly epithelial and predominantly endothelial ROIs. Differential expression analysis between two compartments (airway vs. alveoli vs. blood vessel) was performed using the two-sided non-parametric Wilcoxon test. Correlation analyses were conducted using Spearman coefficients. False discovery rate (FDR) was adjusted using Benjamini– Hochberg correction.

### Xenium spatial transcriptomic analysis of SDC1-4 in mouse embryo

Spatial transcriptomic data were obtained from the publicly available 10x Genomics Xenium Mouse Pup dataset (Xenium Mouse Multi-Tissue Panel, formalin-fixed, paraffin-embedded sagittal section of a developing mouse pup), accessible at: http://www.10xgenomics.com/datasets/mouse-pup-preview-data-xenium-mouse-tissue-atlassing-panel-1-standard (107). Processed Xenium output files were downloaded from the 10x Genomics dataset portal, including transcript coordinates, cell segmentation, cell-by-gene count matrices, and high-resolution tissue morphology images. SDC1 transcript coordinates were mapped across the full section using density heatmaps and single-molecule spatial plots. High-resolution analysis focused on embryonic lung regions, with co-localization assessed using epithelial markers (EpCAM, KRT8) and the endothelial marker VWF. Cell boundaries from segmentation data were overlaid to define cellular context.

### Mouse and human LungMAP (CellRef) reference analysis

Publicly available mouse and human lung single-cell transcriptomic datasets were obtained from the Lung Gene Expression Analysis (LGEA) Web Portal, LungMAP Phase 3, accessible at: https://www.lungmap.net (108–110). All analyses were performed using the LungMAP CellRef reference atlases for human and mouse lung. The LungMAP consortium constructed and curated these reference datasets and used LungMAP CellCards as a unified ontology to harmonize cell identities across datasets. Mouse CellRef includes 95,658 cells generated using Drop-seq across 17 mouse lungs sampled at eight developmental time points from embryonic day 16.5 to postnatal day 28. Human CellRef includes 505,256 cells generated from 104 donor lungs across ten 10x Chromium single-cell RNA-seq datasets. CellRef datasets define 40 mouse and 48 human lung cell types, each annotated using hierarchical lineage structure from LungMAP CellCards (**Supplemental Figures 2, 3**). Major lineages include epithelial, endothelial, mesenchymal, and immune.

### Human bulk tissue and single-cell gene expression of SDC1-4

Human bulk-tissue or single-cell expression of Syndecan family genes (SDC1, SDC2, SDC3, and SDC4) was queried using the GTEx Portal, accessible at https://gtexportal.org (111,112). The GTEx Portal provides uniformly processed bulk RNA sequencing data across multiple non-diseased human tissues, as well as curated single-cell RNA sequencing datasets integrated from publicly available studies. Normalized bulk RNA sequencing expression values (transcripts per million) were extracted for SDC1–4 across tissues and visualized them with violin plots to assess tissue-level expression distributions. Single-cell RNA sequencing data were analyzed across annotated cell populations, and expression patterns of SDC1-4 were compared across cell types to evaluate relative enrichment, particularly between epithelial and endothelial compartments.

### Human gut cell atlas

Single-cell RNA sequencing (scRNA-seq) expression data were obtained from the Human Gut Cell Atlas Portal, accessible at https://www.gutcellatlas.org/pangi.html, which integrates annotated single-cell transcriptomic datasets across multiple regions of the human gastrointestinal tract (113). The resource includes harmonized cell-level expression matrices, curated cell-type annotations, and tissue-of-origin metadata. Log2-normalized transcript expression values were used for downstream visualization. UMAP embeddings provided by the portal were used to visualize global cellular heterogeneity across the atlas. Cell-type annotation included seven major lineage categories based on curated atlas annotations: B and plasma cells, endothelial cells, epithelial cells, mesenchymal cells, myeloid cells, neuronal populations, T cells, and NK cells. Tissue-level annotation encompasses 20 gastrointestinal and associated tissues, including esophagus, stomach, duodenum, jejunum, ileum, colon (ascending, transverse, descending, sigmoid), rectum, appendix, salivary gland, gingival mucosa, buccal mucosa, and mesenteric lymph node (**Supplemental Figure 7**).

### Human brain cell atlas

ScRNA-seq data were obtained from the Human Brain Cell Atlas collection hosted on the CellxGene Portal, accessible at https://cellxgene.cziscience.com/collections/283d65eb-dd53-496d-adb7-7570c7caa443 (114,115). The atlas comprises harmonized, annotated single-cell transcriptomic datasets spanning major regions of the human central nervous system (CNS). Cells were analyzed separately as neuronal and non-neuronal populations based on curated atlas annotations. The neuronal dataset included 2,480,956 cells sampled across 10 CNS regions: cerebellum, cerebral cortex, cerebral nuclei, hippocampal formation, hypothalamus, midbrain, myelencephalon, pons, spinal cord, and thalamic complex. Neurons comprised the majority of cells (2,477,203), with a minor fraction of leukocytes (3,753) identified as infiltrating or resident immune cells. The non-neuronal dataset included 888,263 cells across the same anatomical regions. Major annotated non-neuronal populations included astrocytes, oligodendrocytes, oligodendrocyte precursor cells, CNS macrophages, endothelial cells, pericytes, vascular-associated smooth muscle cells, fibroblasts, ependymal cells, Bergmann glial cells, and choroid plexus epithelial cells (**Supplementary Figure 10**). Normalized gene expression matrices provided by the portal were used for downstream analyses. UMAP embeddings provided by the atlas were used to visualize cellular heterogeneity. Mean expression values were calculated at the cell-type level to evaluate lineage-specific enrichment.

## LIST OF ABBREVIATIONS

CNS: central nervous system
DAPI: 4′,6-diamidino-2-phenylindole
DSP: Digital Spatial Profiling
EOT: Endotheliopathy of trauma
EpCAM: epithelial cell adhesion marker
FDR: False discovery rate
GTEx: Genotype-Tissue Expression Project
KRT8: keratin 8
LGEA: Lung Gene Expression Analysis
NGS: Next Generation Sequencing
OCT: Optimal cutting temperature
PECAM1: platelet endothelial cell adhesion molecule-1
ROIs: Regions of interest
ScRNA-seq: Single-cell RNA-sequencing
SDC1: Syndecan-1
SDC2: Syndecan-2
SDC3: Syndecan-3
SDC4: Syndecan-4
SM: smooth muscle
UCSF: University of California, San Francisco
UMAP: Uniform manifold approximation and projection
VE-cadherin: Vascular endothelial cadherin
VWF: von Willebrand Factor
WTA: Whole Transcriptome Atlas
ZO-1: zonula occludens-1

## AVAILABILITY OF DATA AND MATERIALS

The datasets generated and/or analyzed during the current study are available in the following:

- 10x Genomics Xenium Mouse Pup dataset (Xenium Mouse Multi-Tissue Panel, FFPE sagittal section of a developing mouse pup), http://www.10xgenomics.com/datasets/mouse-pup-preview-data-xenium-mouse-tissue-atlassing-panel-1-standard
- Genotype-Tissue Expression Project Portal, https://gtexportal.org
- Human Gut Cell Atlas Portal, https://www.gutcellatlas.org/pangi.html
- Human Brain Cell Atlas, CellxGene Portal, https://cellxgene.cziscience.com/collections/283d65eb-dd53-496d-adb7-7570c7caa443

All data generated or analyzed during this study are included in this published article and its supplementary information files.

## AUTHOR CONTRIBUTIONS

Conceptualization: SP, HG; Methodology: HG, AT, CK; Data analysis and interpretation: HG, AT, AN, SP; Funding acquisition and supervision: SP; Writing, original draft: AN, HG; Writing, review and editing: AN, HG, SP; Final approval of manuscript: All authors read and approved the final manuscript.

## AI DISCLOSURE

AI (chapGPT, 5.6 Luna) was used solely to proofread the manuscript for wording, grammar, and spelling. AI was not used in the research design, data extraction, analysis, interpretation, or synthesis of the findings. The authors assume full responsibility for ensuring the integrity and accuracy of AI output presented in this article.

## CONFLICT-OF-INTEREST STATEMENT

The authors have declared that no conflict of interest exists.

## FUNDING SUPPORT

This study for supported by supported in part by HDFCCC Laboratory for Cell Analysis Shared Resource Facility through a grant from NIH (P30CA082103).

## Conflict of Interest Statement

The authors have declared that no conflict of interest exists.

## Notes

http://www.10xgenomics.com/datasets/mouse-pup-preview-data-xenium-mouse-tissue-atlassing-panel-1-standard

https://gtexportal.org

https://www.gutcellatlas.org/pangi.html

https://cellxgene.cziscience.com/collections/283d65eb-dd53-496d-adb7-7570c7caa443

### Competing Interest Statement

The authors have declared no competing interest.

## REFERENCES

1. Jaffe EA. Cell biology of endothelial cells. Hum Pathol. 1987 Mar;18(3):234–9.

2. Rahbar E, et al. Endothelial glycocalyx shedding and vascular permeability in severely injured trauma patients. J Transl Med. 2015 Apr 12;13:117.

3. Cheng CK, et al. Biophysical and Biochemical Roles of Shear Stress on Endothelium: A Revisit and New Insights. Circ Res. 2025 Mar 28;136(7):752–772.

4. Neubauer K, Zieger B. Endothelial cells and coagulation. Cell Tissue Res. 2022 Mar;387(3):391–398.

5. Richter RP, et al. Trauma promotes heparan sulfate modifications and cleavage that disrupt homeostatic gene expression in microvascular endothelial cells. Front Cell Dev Biol. 2024 Jul 24;12:1390794.

6. Rehani C, Abdullah S, Kozar RA. Injury induced endotheliopathy: overview, diagnosis, and management. Curr Opin Crit Care. 2025 Jun 1;31(3):237–243.

7. Cardenas JC, Dong JF, Kozar RA. Injury-induced endotheliopathy: What you need to know. J Trauma Acute Care Surg. 2023 Oct 1;95(4):454–463.

8. Rehani C, et al. A Component Analysis of Endotheliopathy of Trauma. Res Sq [Preprint]. 2026 May 19:rs.3.rs-9318157.

9. Wu F, et al. Resuscitative Strategies to Modulate the Endotheliopathy of Trauma: From Cell to Patient. Shock. 2020 May;53(5):575–584.

10. Naumann DN, et al. Endotheliopathy of Trauma is an on-Scene Phenomenon, and is Associated with Multiple Organ Dysfunction Syndrome: A Prospective Observational Study. Shock. 2018 Apr;49(4):420–428.

11. Holcomb JB, Pati S. Optimal trauma resuscitation with plasma as the primary resuscitative fluid: the surgeon’s perspective. Hematology Am Soc Hematol Educ Program. 2013;2013:656–9.

12. White NJ, et al. Hemorrhagic blood failure: Oxygen debt, coagulopathy, and endothelial damage. J Trauma Acute Care Surg. 2017 Jun;82(6S Suppl 1):S41–S49.

13. Moore KH, Murphy HA, George EM. The glycocalyx: a central regulator of vascular function. Am J Physiol Regul Integr Comp Physiol. 2021 Apr 1;320(4):R508–R518.

14. Gaudette S, Hughes D, Boller M. The endothelial glycocalyx: Structure and function in health and critical illness. J Vet Emerg Crit Care (San Antonio).2020 Mar;30(2):117–134.

15. Hahn RG, Patel V, Dull RO. Human glycocalyx shedding: Systematic review and critical appraisal. Acta Anaesthesiol Scand. 2021 May;65(5):590–606.

16. Uchimido R, Schmidt EP, Shapiro NI. The glycocalyx: a novel diagnostic and therapeutic target in sepsis. Crit Care. 2019 Jan 17;23(1):16.

17. Becker BF, et al. Degradation of the endothelial glycocalyx in clinical settings: searching for the sheddases. Br J Clin Pharmacol. 2015 Sep;80(3):389–402.

18. Diebel LN, et al. Acute hyperglycemia exacerbates trauma-induced endothelial and glycocalyx injury: An in vitro model. J Trauma Acute Care Surg. 2018 Nov;85(5):960–967.

19. Johansson PI, Stensballe J, Ostrowski SR. Shock induced endotheliopathy (SHINE) in acute critical illness - a unifying pathophysiologic mechanism. Crit Care. 2017 Feb 9;21(1):25.

20. Carge MJ, Liberati DM, Diebel LN. A biomimetic shock model on the effect of endothelial aging on vascular barrier properties. J Trauma Acute Care Surg. 2021 Nov 1;91(5):849–855.

21. Martin JV, Liberati DM, Diebel LN. Disparate effects of catecholamines under stress conditions on endothelial glycocalyx injury: An in vitro model. Am J Surg. 2017 Dec;214(6):1166–1172.

22. Braunstein M, et al. Effect on Syndecan-1 and Hyaluronan Levels Depending on Multiple Organ Failure, Coagulopathy and Survival: An Observational Study in Major Trauma Patients. J Clin Med. 2024 Nov 10;13(22):6768.

23. Inoda A, et al. Glycocalyx shedding as a clinical biomarker in critical illness. Exp Mol Pathol. 2025 Dec;144:104997.

24. Matsumoto H, et al. Association of endotheliopathy with coagulofibrinolytic reactions and disseminated intravascular coagulation after trauma: a retrospective observational study. Sci Rep. 2024 Nov 28;14(1):29630.

25. Wade CE, et al. Absences of Endothelial Microvesicle Changes in the Presence of the Endotheliopathy of Trauma. Shock. 2019 Feb;51(2):180–184.

26. Kozar RA, et al. Plasma restoration of endothelial glycocalyx in a rodent model of hemorrhagic shock. Anesth Analg. 2011 Jun;112(6):1289–95.

27. Rangarajan S, et al. Heparanase-enhanced Shedding of Syndecan-1 and Its Role in Driving Disease Pathogenesis and Progression. J Histochem Cytochem. 2020 Dec;68(12):823–840.

28. Baljinnyam T, et al. High molecular weight sodium hyaluronate improves survival of syndecan-1-deficient septic mice by inhibiting neutrophil migration. PLoS One. 2021 Apr 30;16(4):e0250327.

29. Tarbell JM, Ebong EE. The endothelial glycocalyx: a mechano-sensor and -transducer. Sci Signal. 2008 Oct 7;1(40):pt8.

30. Curry FE, Adamson RH. Endothelial glycocalyx: permeability barrier and mechanosensor. Ann Biomed Eng. 2012 Apr;40(4):828–39.

31. Weinbaum S, et al. Mechanotransduction and flow across the endothelial glycocalyx. Proc Natl Acad Sci U S A. 2003 Jun 24;100(13):7988–95.

32. Richter RP, et al. Plasma Angiopoietin-2/-1 Ratio is Elevated and Angiopoietin-2 Levels Correlate With Plasma Syndecan-1 Following Pediatric Trauma. Shock. 2019 Sep;52(3):340–346.

33. Hayashida K, Parks WC, Park PW. Syndecan-1 shedding facilitates the resolution of neutrophilic inflammation by removing sequestered CXC chemokines. Blood. 2009 Oct 1;114(14):3033–43.

34. Dujardin RWG, et al. Shock-Driven Endotheliopathy in Trauma Patients Is Associated with Leucocyte Derived Extracellular Vesicles. Int J Mol Sci. 2022 Dec 15;23(24):15990.

35. Baucom MR, et al. Syndecan-1 as the Effect or Effector of the Endothelial Inflammatory Response? J Surg Res. 2024 Mar;295:611–618.

36. Ostrowski SR, Johansson PI. Endothelial glycocalyx degradation induces endogenous heparinization in patients with severe injury and early traumatic coagulopathy. J Trauma Acute Care Surg. 2012 Jul;73(1):60–6.

37. Straat M, et al. Effect of transfusion of fresh frozen plasma on parameters of endothelial condition and inflammatory status in non-bleeding critically ill patients: a prospective substudy of a randomized trial. Crit Care. 2015 Apr 15;19(1):163.

38. Yang Z, et al. Traumatized triad of complementopathy, endotheliopathy, and coagulopathy - Impact on clinical outcomes in severe polytrauma patients. Front Immunol. 2022 Oct 20;13:991048.

39. Krocker JD, et al. Exploratory Investigation of the Plasma Proteome Associated with the Endotheliopathy of Trauma. Int J Mol Sci. 2022 Jun 1;23(11):6213.

40. Ban K, et al. Plasma-Mediated Gut Protection After Hemorrhagic Shock is Lessened in Syndecan-1- /- Mice. Shock. 2015 Nov;44(5):452–7.

41. Johansson PI, et al. A high admission syndecan-1 level, a marker of endothelial glycocalyx degradation, is associated with inflammation, protein C depletion, fibrinolysis, and increased mortality in trauma patients. Ann Surg. 2011 Aug;254(2):194–200.

42. Lopez E, et al. Antithrombin III Contributes to the Protective Effects of Fresh Frozen Plasma Following Hemorrhagic Shock by Preventing Syndecan-1 Shedding and Endothelial Barrier Disruption. Shock. 2020 Feb;53(2):156–163.

43. Kozar RA, Pati S. Syndecan-1 restitution by plasma after hemorrhagic shock. J Trauma Acute Care Surg. 2015 Jun;78(6 Suppl 1):S83-6.

44. Nelson A, et al. Circulating syndecans during critical illness. APMIS. 2017 May;125(5):468–475.

45. Wu F, et al. Loss of Syndecan-1 Abrogates the Pulmonary Protective Phenotype Induced by Plasma After Hemorrhagic Shock. Shock. 2017 Sep;48(3):340–345.

46. Peng Z, et al. Syndecan 1 plays a novel role in enteral glutamine’s gut-protective effects of the postischemic gut. Shock. 2012 Jul;38(1):57–62.

47. Wu F, Kozar RA. Fibrinogen Protects Against Barrier Dysfunction Through Maintaining Cell Surface Syndecan-1 In Vitro. Shock. 2019 Jun;51(6):740–744.

48. Haywood-Watson RJ, et al. Modulation of syndecan-1 shedding after hemorrhagic shock and resuscitation. PLoS One. 2011;6(8):e23530.

49. Halbgebauer R, et al. Hemorrhagic shock drives glycocalyx, barrier and organ dysfunction early after polytrauma. J Crit Care. 2018 Apr;44:229–237.

50. Gonzalez Rodriguez E, et al. Traumatic brain injury is associated with increased syndecan-1 shedding in severely injured patients. Scand J Trauma Resusc Emerg Med. 2018 Nov 21;26(1):102.

51. Xie WW, et al. Ding YJ, Bhandari S, Li H, Chen HS, Jin SW, Weng HX, Hao Y. Clinical value of syndecan-1 levels in trauma brain injury: A meta-analysis. Shock. 2024. Jan 1;61(1):49–54.

52. Wallen TE, et al. Defining Endotheliopathy in Murine Polytrauma Models. Shock. 2022 Jun 1;57(6):291–298.

53. Zeineddin A, et al. Biomarkers of endothelial cell dysfunction persist beyond resuscitation in patients with hemorrhagic shock. J Trauma Acute Care Surg. 2022 Nov 1;93(5):572–578.

54. Albert V, et al. Acute Traumatic Endotheliopathy in Isolated Severe Brain Injury and Its Impact on Clinical Outcome. Med Sci (Basel). 2018 Jan 16;6(1):5.

55. Hofmann N, et al. Experimental Models of Endotheliopathy: Impact of Shock Severity. Shock. 2018 May;49(5):564–571.

56. Rodriguez EG, et al. Syndecan-1: A Q;uantitative Marker for the Endotheliopathy of Trauma. J Am Coll Surg. 2017 Sep;225(3):419–427.

57. Johansson PI, et al. Traumatic Endotheliopathy: A Prospective Observational Study of 424 Severely Injured Patients. Ann Surg. 2017 Mar;265(3):597–603.

58. Gruen DS, et al. PAMPer study group: Prehospital plasma is associated with distinct biomarker expression following injury. JCI Insight. 2020 Apr 23;5(8):e135350.

59. Suzuki K, et al. Syndecan-1 as a severity biomarker for patients with trauma. Front Med (Lausanne). 2022 Sep 27;9:985955.

60. Morgan KM, et al. Endotheliopathy of trauma in children: The association of syndecan-1 with injury and poor outcomes. J Trauma Acute Care Surg. 2024 Apr 1;96(4):566–572.

61. Wei S, et al. Elevated Syndecan-1 after Trauma and Risk of Sepsis: A Secondary Analysis of Patients from the Pragmatic, Randomized Optimal Platelet and Plasma Ratios (PROPPR) Trial. J Am Coll Surg. 2018 Dec;227(6):587–595.

62. Gruen DS, et al. Prehospital tranexamic acid is associated with a dose-dependent decrease in syndecan-1 after trauma: A secondary analysis of a prospective randomized trial. J Trauma Acute Care Surg. 2023 Nov 1;95(5):642–648.

63. Anderson TN, et al. Early Tranexamic Acid Administration After Traumatic Brain Injury Is Associated With Reduced Syndecan-1 and Angiopoietin-2 in Patients With Traumatic Intracranial Hemorrhage. J Head Trauma Rehabil. 2020 Sep/Oct;35(5):317–323.

64. Peng Z, et al. Fresh frozen plasma lessens pulmonary endothelial inflammation and hyperpermeability after hemorrhagic shock and is associated with loss of syndecan 1. Shock. 2013 Sep;40(3):195–202.

65. Anand T, et al. Decreased Glycocalyx Shedding on Presentation in Hemorrhaging Geriatric Trauma Patients. J Surg Res. 2024 Jan;293:709–716.

66. Kregel HR, et al. Shock-Induced Endothelial Dysfunction is Present in Patients With Occult Hypoperfusion After Trauma. Shock. 2022 Jan 1;57(1):106–112.

67. Hatton GE, et al. Endothelial Dysfunction is Associated With Increased Incidence, Worsened Severity, and Prolonged Duration of Acute Kidney Injury After Severe Trauma. Shock. 2021 Mar 1;55(3):311–315.

68. Baucom MR, et al. Blood component resuscitative strategies to mitigate endotheliopathy in a murine hemorrhagic shock model. J Trauma Acute Care Surg. 2023 Jul 1;95(1):21–29.

69. Chipman AM, et al. Fresh frozen plasma attenuates lung injury in a novel model of prolonged hypotensive resuscitation. J Trauma Acute Care Surg. 2020 Aug;89(2S Suppl 2):S118–S125.

70. Weinberg L, et al. Trajectory of plasma syndecan-1 and heparan sulphate during major surgery: A retrospective observational study. Acta Anaesthesiol Scand. 2023 Jan;67(1):4–11.

71. Keyloun JW, et al. Circulating Syndecan-1 and Tissue Factor Pathway Inhibitor, Biomarkers of Endothelial Dysfunction, Predict Mortality in Burn Patients. Shock. 2021 Aug 1;56(2):237–244.

72. Bonson G, et al. Syndecan-1 Levels in Trauma and Burn Patients Remain Elevated During Resuscitation and Correlate With Coagulopathy. J Am Coll Emerg Physicians Open.2026 Apr 7;7(3):100350.

73. Ogawa F, et al. Temporal change in Syndecan-1 as a therapeutic target and a biomarker for the severity classification of COVID-19. Thromb J. 2021 Aug 16;19(1):55.

74. Dixon A, et al. Acute respiratory distress syndrome, acute kidney injury, and mortality after trauma are associated with increased circulation of syndecan-1, soluble thrombomodulin, and receptor for advanced glycation end products. J Trauma Acute Care Surg. 2024 Feb 1;96(2):319–325.

75. Piotti A, et al. Endothelial damage in septic shock patients as evidenced by circulating syndecan-1, sphingosine-1-phosphate and soluble VE-cadherin: a substudy of ALBIOS. Crit Care. 2021 Mar 19;25(1):113.

76. Kelly EJ, et al. Endothelial damage occurs early after inhalation injury as measured by increased syndecan-1 levels. J Burn Care Res. 2023 Jul 5;44(4):769–774.

77. Vigiola, et al. Plasma Ameliorates Endothelial Dysfunction in Burn Injury. J Surg Res. 2019 Jan;233:459–466.

78. Welling H, et al. Endothelial glycocalyx shedding in patients with burns. Burns. 2020 Mar;46(2):386–393.

79. Eglin CM, et al. Plasma biomarkers of endothelial function, inflammation and oxidative stress in individuals with non-freezing cold injury. Exp Physiol. 2023 Mar;108(3):448–464.

80. Meneses GC, et al. Endothelial biomarkers as predictors for haemodialysis need in severe leptospirosis patients (Weil’s disease). Trop Med Int Health. 2022 Aug;27(8):727–734.

81. Ziedins EE, et al. Markers of Endothelial Injury in Extracorporeal Membrane Oxygenation: A New Risk Assessment Method. J Surg Res. 2024 Dec;304:391–399.

82. Kelly EJ, et al. Endothelial dysfunction is dampened by early administration of fresh frozen plasma in a rodent burn shock model. J Trauma Acute Care Surg. 2024 Oct 1;97(4):520–528.

83. Johansson PI, et al. Association between sympathoadrenal activation, fibrinolysis, and endothelial damage in septic patients: a prospective study. J Crit Care. 2014 Jun;29(3):327–33.

84. Zhang Q, et al. Longitudinal Assessment of Plasma Syndecan-1 Predicts 60-Day Mortality in Patients with COVID-19. J Clin Med. 2023 Jan 10;12(2):552.

85. Xu D, et al. Dong C, Zhao Z, Hou S. Syndecan-1 as a biomarker of endothelial glycocalyx injury in sepsis: a meta-analysis of associations with shock, organ complications and mortality. Postgrad Med J. 2026 Apr 22:qgag053.

86. Stensballe J, et al. Resuscitation of Endotheliopathy and Bleeding in Thoracic Aortic Dissections: The VIPER-OCTA Randomized Clinical Pilot Trial. Anesth Analg. 2018 Oct;127(4):920–927.

87. Pusateri AE, et al. Fibrinolytic dysfunction and endotheliopathy after major thermal injury: Considerations needed for new approaches to burn shock resuscitation. Shock. 2025 Jan 1;63(1):11–18.

88. Uchida K, et al. Glycocalyx biomarkers as early predictors of endotheliopathy in pediatric and young adult hematopoietic stem cell transplantation patients. Front Oncol. 2026 May 8;16:1789000.

89. Bell JD, et al. Biomarkers of Glycocalyx Injury are Associated with Delayed Cerebral Ischemia Following Aneurysmal Subarachnoid Hemorrhage: A Case Series Supporting a New Hypothesis. Neurocrit Care. 2017 Jun;26(3):339–347.

90. Brauer R, et al. Syndecan-1 Attenuates Lung Injury during Influenza Infection by Potentiating c-Met Signaling to Suppress Epithelial Apoptosis. Am J Respir Crit Care Med. 2016 Aug 1;194(3):333–44.

91. Palaiologou M, Delladetsima I, Tiniakos D. CD138 (syndecan-1) expression in health and disease. Histol Histopathol. 2014 Feb;29(2):177–89.

92. Diab L, Al Kattar S, Oueini N, Hawi J, Chrabieh A, Dosh L, Jurjus R, Leone A, Jurjus A. Syndecan-1: a key player in health and disease. Immunogenetics. 2024 Dec 17;77(1):9.

93. Schønemann-Lund M, et al. Novel subgroups in acute respiratory failure based on the trajectories of three endotheliopathy biomarkers: A cohort study. Acta Anaesthesiol Scand. 2023 Aug;67(7):896–908.

94. Thanapaul RJRS, et al. Pathophysiology-guided biomarkers and therapeutics for precision trauma medicine in polytrauma with musculoskeletal injuries. Mil Med Res. 2026 Dec;13(1):100014.

95. Contreras HR, et al. The expression of syndecan-1 and -2 is associated with Gleason score and epithelial-mesenchymal transition markers, E-cadherin and beta-catenin, in prostate cancer. Urol Oncol. 2010 Sep-Oct;28(5):534–40.

96. Zhang Y, et al. Cell surface-anchored syndecan-1 ameliorates intestinal inflammation and neutrophil transmigration in ulcerative colitis. J Cell Mol Med. 2017 Jan;21(1):13–25.

97. Alaeddini M, Yazdani F, Etemad-Moghadam S. Stromal and epithelial syndecan-1 expression in benign and malignant salivary gland tumors: which is more reflective of behavior? Braz J Otorhinolaryngol. 2021 Mar-Apr;87(2):171–177.

98. Nam EJ, et al. Age-Related Variation of Syndecan-1 Levels in Saliva and Plasma of Healthy Individuals. Clin Lab. 2021 Sep 1;67(9).

99. Randolph GJ, et al. The Lymphatic System: Integral Roles in Immunity. Annu Rev Immunol. 2017 Apr 26;35:31–52.

100. Day RM, et al. Regulation of epithelial syndecan-1 expression by inflammatory cytokines. Cytokine. 2003 Mar 7;21(5):224–33.

101. Guo M, et al. Syndecan-1 shedding destroys epithelial adherens junctions through STAT3 after renal ischemia/reperfusion injury. iScience. 2023 Oct 14;26(11):108211.

102. Yao C, et al. Syndecan-1 Promotes Alveolar Type 2 Epithelial Cell Senescence during Lung Fibrosis. bioRxiv [Preprint]. 2026 Apr 1:2026.03.16.712248.

103. Celie JW, et al. Tubular epithelial syndecan-1 maintains renal function in murine ischemia/reperfusion and human transplantation. Kidney Int. 2012 Apr;81(7):651–61.

104. Lundin M, et al. Epithelial syndecan-1 expression is associated with stage and grade in colorectal cancer. Oncology. 2005;68(4-6):306–13.

105. Ahmed Haji Omar A, et al. Epithelial and stromal syndecan-1 and -2 are distinctly expressed in oral- and cutaneous squamous cell carcinomas. J Oral Pathol Med. 2013 May;42(5):389–95.

106. Barry M, et al. Mesenchymal stem cell extracellular vesicles mitigate vascular permeability and injury in the small intestine and lung in a mouse model of hemorrhagic shock and trauma. J Trauma Acute Care Surg. 2022 Mar 1;92(3):489–498.

107. 10x Genomics. Whole Mouse Pup Preview Data (Xenium Mouse Tissue Atlassing Panel). http://www.10xgenomics.com/datasets/mouse-pup-preview-data-xenium-mouse-tissue-atlassing-panel-1-standard. Published August 8, 2023. Accessed August 6, 2026.

108. Du Y, et al. Lung Gene Expression Analysis (LGEA): An integrative web portal for comprehensive gene expression analysis in lung development and disease. Am J Respir Cell Mol Biol. 2021;64(3):285–9.

109. Guo M, et al. Guided construction of single cell reference for human and mouse lung. Nat Commun. 2023 Jul 29;14(1):4566.

110. LungMap. Lung Gene Expression Analysis (LGEA) Web Portal. https://www.lungmap.net. Published 2023. Accessed August 6, 2026.

111. Liu S, Gao Y, Canela-Xandri O, Wang S, Yu Y, Cai W, GTEx Consortium. The Genotype-Tissue Expression (GTEx) pilot analysis: multi-tissue gene regulation in humans. Nat Genet. 2013;45(5):580–585.

112. Genotype-Tissue Expression Project (GTEx) Portal. https://gtexportal.org. Published 2026. Accessed August 6, 2026.

113. Gut Cell Survey. Human Gut Cell Atlas Portal. https://www.gutcellatlas.org/pangi.html. Published 2026. Accessed August 6, 2026.

114. Siletti K, et al. Transcriptomic diversity of cell types across the adult human brain. Science. 2023 Oct 13;382(6667):eadd7046.

115. Chan Zuckerberg Initiative, CellxGene Portal. Human Brain Cell Atlas v1.0. https://cellxgene.cziscience.com/collections/283d65eb-dd53-496d-adb7-7570c7caa443. Published October 13, 2023. Accessed August 6, 2026.

