## Supplemental Figures for "Rethinking Syndecan-1 as a Marker of Trauma-Induced Endothelial Injury"

**SUPPLEMENTAL FIGURE LEGENDS:**

**Supplemental Figure 1: Overview of the GeoMx Digital Spatial Profiling (DSP) workflow. Day 1**: Fresh-frozen lung lobes were sectioned and stained with morphology markers and incubated with whole transcriptome atlas mouse probes overnight. **Day 2**: Slides were placed on GeoMx station and processed for region of interest (ROI) selection. These ROIs were collected in 96-well plates. **Day 3**: Released oligos undergo library preparation followed by next-generation sequencing. Sequencing output files (FastQ) are processed and converted for DSP data analysis, including quality control, spatial segmentation, and expression quantification.

**Supplemental Figure 2: Mouse lung map cell types. (A)** A comprehensive cell-type reference (CellRef) for mouse lung tissues, grouped into four major cell lineages: endothelial, epithelial, mesenchymal, and immune cells. CellRef consolidates single-cell transcriptomic datasets of 17 mouse lung samples to construct the Lung MAP single-cell reference, defining 40 distinct mouse lung cell types. **(B)** Each cell type is color-coded to illustrate the full spectrum of mouse lung cell populations used in **Figure 5**. The single-cell RNA sequencing data were obtained through the Lung MAP Lung Gene Expression Analysis (LGEA) web portal (https://www.lungmap.net).

**Supplemental Figure 3: Human lung map cell types. (A)** A comprehensive cell-type reference (CellRef) for human lung tissues, grouped into four major cell lineages: endothelial, epithelial, mesenchymal, and immune cells. CellRef consolidated single-cell transcriptomic datasets of 104 normal human lungs to construct the Lung MAP single-cell reference, defining 48 human lung cell types. **(B)** Each cell type is color-coded to illustrate the full spectrum of human lung cell populations used in **Figure 6**. The single-cell RNA-seq data were obtained through the Lung MAP Lung Gene Expression Analysis (LGEA) web portal (https://www.lungmap.net).

**Supplemental Figure 4: Human tissue gene expression for syndecan-1 (SDC1). (A)** Bulk RNA sequencing expression levels of SDC1 are shown across different human tissues profiled by the Genotype-Tissue Expression (GTEx) Project. Violin plots illustrate the distribution of normalized SDC1 transcript abundance within each tissue. (**B**) Violin plots depict the distribution of SDC1 expression at the single-cell level across diverse human tissues and cell types. Bulk and single-cell RNA-seq expression data were obtained from the GTEx Portal (https://www.gtexportal.org).

**Supplemental Figure 5: Bulk RNA sequencing expression profiles of syndecan-2, syndecan-3, and syndecan-4 (SDC2, SDC3, SDC4) across human tissues.** Bulk RNA sequencing expression levels of **(A)** SDC2, **(B)** SDC3, and **(C)** SDC4 are shown across different human tissues profiled by the Genotype-Tissue Expression (GTEx) Project. Violin plots illustrate the distribution of normalized SDC2, SDC3, and SDC4 transcript abundance within each tissue, capturing inter-individual variability and tissue-specific expression patterns. Data were obtained from the GTEx Portal (https://www.gtexportal.org).

**Supplemental Figure 6: Single-cell expression profiles of SDC2, SDC3, and SDC4 across human tissues and cell types.** Violin plots depict the distribution of **(A)** SDC2, **(B)** SDC3, and **(C)** SDC4 expression at the single-cell level across diverse human tissues and cell types profiled by the Genotype-Tissue Expression (GTEx) Project. Single-cell expression data were obtained from the GTEx Portal (https://www.gtexportal.org).

**Supplemental Figure 7: Human gut cell atlas annotation.** Summary of the tissue and cell-type composition of the human multiorgan single-cell gut atlas used in **Supplemental Figures 8 and 9**. The table summarizes the distribution of 1,077,241 single cells across 20 major human gut and gut-associated tissues and 7 major cellular lineages, including B and plasma cells (149,240 cells), endothelial cells (52,804), epithelial cells (371,724), mesenchymal cells (270,956), myeloid cells (33,213), neural populations (23,121), and T/NK cells (176,183). Single-cell expression data were obtained from the Human Gut Cell Atlas Portal (https://www.gutcellatlas.org/pangi.html).

**Supplemental Figure 8: SDC1 expression patterns in the human gut single-cell atlas. (A)** Uniform manifold approximation and projection (UMAP) embeddings display log2-normalized transcript levels of SDC1 across all cells in the human gut single-cell atlas. Warmer colors denote higher transcript levels, highlighting selective enrichment in epithelial and plasma cell compartments. **(B)** UMAP visualization colored by 7 major cell-type lineages: B & plasma cells, endothelial cells, epithelial cells, mesenchymal cells, myeloid cells, neuronal populations, and T/NK cells. **(C)** UMAP visualization colored by 20 human tissues. Single-cell expression data were obtained from the Human Gut Cell Atlas Portal (https://www.gutcellatlas.org/pangi.html). **(D)** Table summarizing SDC1 expression by cell type, including the number of cells for each cell type, proportion of SDC1-expressing cells, and the mean SDC1 expression. Strongest SDC1 expression is observed in B/plasma cells and epithelial cells, while endothelial cells show minimal expression of SDC1.

**Supplemental Figure 9: UMAP visualization of SDC2, SDC3, and SDC4 expression in different cell types and tissue origins in the human gut single-cell atlas.** UMAP embeddings display log2-normalized transcript levels of **(A)** SDC2, **(B)** SDC3, and **(C)** SDC4 across all cells in the atlas. Warmer colors indicate higher expression levels. UMAP visualization of 7 major cell-type lineages and 20 human tissues can be seen in **Supplemental Figure 8B** and **8C**, respectively. Single-cell expression data were obtained from the Human Gut Cell Atlas Portal (https://www.gutcellatlas.org/pangi.html).

**Supplemental Figure 10: Human brain cell atlas annotation.** Summary of neuronal and non-neuronal cell composition across human central nervous system (CNS) regions used in **Supplemental Figures 11 and 12**. **(A)** Neuronal cell distribution across CNS anatomical regions. The table summarizes 2,480,956 neuronal cells sampled from 10 major brain and spinal cord regions, including cerebellum, cerebral cortex, cerebral nuclei, hippocampal formation, hypothalamus, midbrain, myelencephalon, pons, spinal cord, and thalamic complex. Across all regions, neurons account for most cells (2,477,203), with a small fraction of leukocytes (3,753) detected as infiltrating or resident immune cells. **(B)** Non-neuronal cell distribution across CNS regions. The table summarizes 888,263 non-neuronal cells across the same 10 major brain and spinal cord regions. Major non-neuronal cell classes include Bergmann glial cells (8,041), astrocytes (155,025), CNS macrophages (91,838), choroid plexus epithelial cells (7,689), endothelial cells (5,165), ependymal cells (5,882), fibroblasts (9,156), oligodendrocytes (494,966), oligodendrocyte precursor cells (39,733), pericytes (3,963), and vascular-associated smooth muscle cells (1,074). Single-cell expression data were obtained from the Human Brain Cell Atlas Portal (https://cellxgene.cziscience.com/collections/283d65eb-dd53-496d-adb7-7570c7caa443).

**Supplemental Figure 11: SDC1 expression across neuronal and non-neuronal cell populations in the human brain single-cell atlas. (A-C)** SDC1 expression in neurons. **(A)** UMAP visualization of neuron-only single-cell data, with cells colored by normalized SDC1 expression. Neuronal populations show negligible SDC1 signal across all clusters. **(B)** Neuronal cells are colored by brain-region origin, including cerebral cortex, pons, midbrain, hypothalamus, cerebellum, hippocampal formation, myelencephalon, spinal cord, thalamic complex, and cerebral nuclei, demonstrating broad anatomical sampling across the CNS. **(C)** The mean expression level of SDC1 in each cell type. **(D-G)** SDC1 expression in non-neuronal cells. **(D)** UMAP visualization of SDC1 expression in non-neuronal single-cell data. **(E)** Non-neuronal cells are colored by anatomical origin, matching the regions represented in the neuronal dataset in (**B**). **(F)** UMAP colored by annotated non-neuronal identities, including fibroblasts, ependymal cells, endothelial cells, astrocytes, oligodendrocytes, oligodendrocyte precursor cells (OPCs), Bergmann glia, pericytes, vascular smooth muscle cells, choroid plexus epithelial cells, and CNS macrophages. **(G)** Summary of SDC1 mean expression within each cell group, confirming uniformly low to absent expression in non-neuronal lineages.

**Supplemental Figure 12: UMAP visualization of syndecan-2, syndecan-3, and syndecan-4 (SDC2, SDC3, SDC4) expression across neuronal and non-neuronal cell populations in the human brain single-cell atlas. (A-B)** SDC2, **(C-D)** SDC4, and **(E-F)** SDC3 expression in neurons and non-neuronal cells. UMAP visualizations show SDC2, SDC3, and SDC4 transcript levels across neuronal (**A**, **C, E**) and non-neuronal (**B**, **D, F**) subsets of the human brain single-cell atlas. SDC2 and SDC4 expressions are nearly absent in both compartments, with no detectable enrichment in any neuron, and non-neuron cell subtype. SDC3 shows low but detectable expression in subclusters of neuronal and non-neuronal cells. (**G-H**) Bar plots depict the distribution of SDC3 expression across all CNS cells, showing overall low-level SDC3 expression in neuronal cells (**G**), and across different non-neuronal cell types (**H**). Single-cell expression data were obtained from the Human Brain Cell Atlas Portal (https://cellxgene.cziscience.com/collections/283d65eb-dd53-496d-adb7-7570c7caa443).

### Supplemental Figure 1.

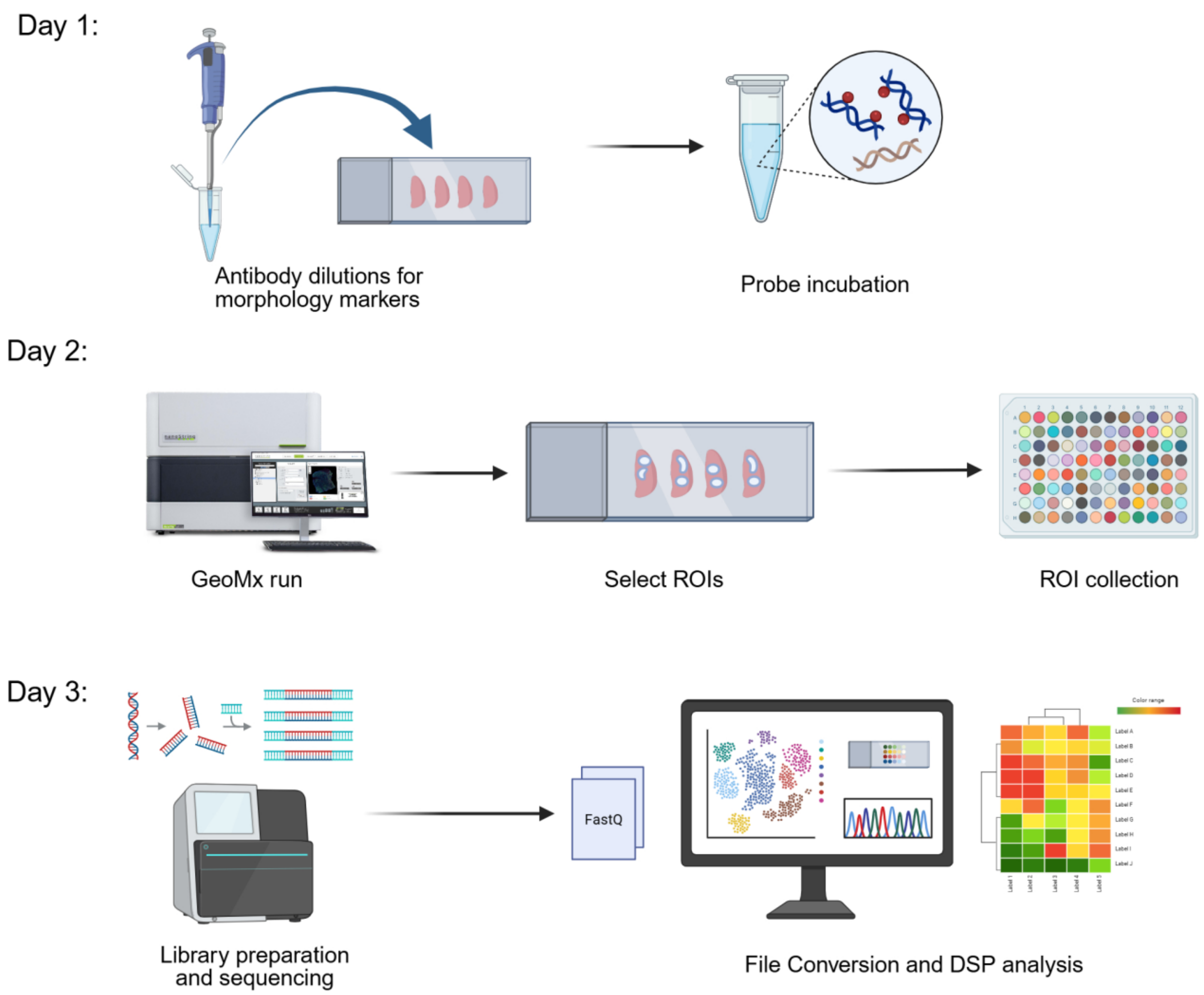

### Supplemental Figure 2.

#### A. CellRef: mouse cell types (40)

| Endothelial Cells | Cell Type |
|---|---|
| AEC | Arterial endothelial cell (subtype of Vacular endothelial cell) |
| CAP1 | Capillary cell 1 (subtype of Vascular endothelial cell) |
| CAP2 | Capillary cell 2 (subtype of Vascular endothelial cell) |
| EPC | Endothelial Progenitor Cell (subtype of Endothelium) |
| LEC | Lymphatic endothelial cell (subtype of Endothelial cell) |
| VEC | Venous endothelial cell (subtype of Vascular endothelial cell) |
| **Epithelial Cells** | **Cell Type** |
| AT1 | Alveolar type 1 cell (subtype of Distal epithelial Cell) |
| AT1/AT2 | Alveolar type 1/2 cell (subtype of Distal epithelial Cell) |
| AT2 | Alveolar type 2 cell (subtype of Distal epithelial Cell) |
| Basal | Basal cell (subtype of Airway epithelial Cell) |
| Ciliated | Ciliated cell (subtype of Airway epithelial Cell) |
| Deuterosomal | Deuterosomal Cell (subtype of Epithelium) |
| PNEC | Pulmonary neuroendocrine cell (subtype of Airway epithelial Cell) |
| Secretory | Secretory cell (subtype of Airway epithelial Cell) |
| Sox9_Epi | Sox9+/Id2+ Epithelial Cell (subtype of Epithelium) |
| **Mesenchymal Cells** | **Cell Type** |
| AF1 | Alveolar fibroblast 1 (subtype of Matrix Fibroblast) |
| AF2 | Alveolar fibroblast 2 (subtype of Matrix Fibroblast) |
| ASMC | Airway Smooth Muscle (subtype of Mesenchymal cell) |
| Chondrocyte | Chondrocytes (subtype of Mesenchyme) |
| Mesothelial | Mesothelium (subtype of Mesenchyme) |
| Pericyte | Pericyte (subtype of Mesenchymal Cell) |
| PMP | Proliferative mesenchymal progenitor (subtype of Mesenchymal Cell) |
| SCMF | Secondary crest myofibroblast cell (subtype of Mesenchymal Cell) |
| VSMC | Vascular smooth muscle (subtype of Mesenchymal Cell) |
| **Immune Cells** | **Cell Type** |
| AM | Alveolar macrophage (subtype of Myeloid) |
| B | B Lymphocyte (subtype of Lymphocyte) |
| Basophils | Basophil (subtype of Myeloid) |
| CD4_T | CD4+ T Cell (subtype of Immune) |
| CD8_T | CD8+ T Cell (subtype of Immune) |
| cDC1 | Classical DC subset 1 (subtype of Dendritic cell) |
| cDC2 | Classical DC subset 2 (subtype of Dendritic cell) |
| ILC | Innate lymphoid cells (subtype of Lymphocyte) |
| IM | Interstitial macrophage (subtype of Myeloid) |
| iMON | Inflammatory monocytes (subtype of Monocyte) |
| maDC | maDC, a highly specific marker of mature Dendritic cells (subtype of Myeloid) |
| Mast | Mast (subtype of Myeloid) |
| Neutrophil | Neutrophil (subtype of Myeloid) |
| NK | Natural Killer Cell (subtype of Lymphocyte) |
| Platelet | Megakaryocyte/Platelet (subtype of Immune Cell) |
| Treg | T Regulatory Cell (subtype of Immune) |

#### B. Color-coded mouse lung cell types

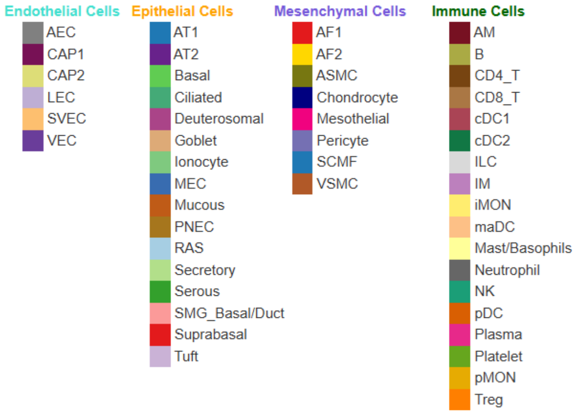

### Supplemental Figure 3.

#### A. CellRef: human cell types (48)

| Endothelial Cells | Cell Type |
|---|---|
| AEC | Arterial endothelial cell (subtype of Vacular endothelial cell) |
| CAP1 | Capillary cell 1 (subtype of Vascular endothelial cell) |
| CAP2 | Capillary cell 2 (subtype of Vascular endothelial cell) |
| LEC | Lymphatic endothelial cell (subtype of Endothelial cell) |
| SVEC | Systemic Venous Endothelial Cell (subtype of Endothelium) |
| VEC | Venous endothelial cell (subtype of Vascular endothelial cell) |
| **Epithelial Cells** | **Cell Type** |
| AT1 | Alveolar type 1 cell (subtype of Distal epithelial Cell) |
| AT2 | Alveolar type 2 cell (subtype of Distal epithelial Cell) |
| Basal | Basal cell (subtype of Airway epithelial Cell) |
| Ciliated | Ciliated cell (subtype of Airway epithelial Cell) |
| Deuterosomal | Deuterosomal Cell (subtype of Epithelium) |
| Goblet | Goblet cell (subtype of Airway epithelial Cell) |
| Ionocyte | Ionocyte cell (subtype of Airway epithelial Cell) |
| MEC | Myoepithelial cell (subtype of Airway epithelial Cell) |
| Mucous | Mucous cell (subtype of Epithelial Cell) |
| PNEC | Pulmonary neuroendocrine cell (subtype of Airway epithelial Cell) |
| RAS | Respiratory Airway Secretory Cells (subtype of Epithelium) |
| Secretory | Secretory cell (subtype of Airway epithelial Cell) |
| Serous | Serous cell (subtype of Airway epithelial Cell) |
| SMG_Basal/Duct | Terminal ciliated duct basal cells (subtype of Epithelial cell) |
| Suprabasal | Suprabasal Cell (subtype of Epithelium) |
| Tuft | Tuft cell (subtype of Airway epithelial Cell) |
| **Mesenchymal Cells** | **Cell Type** |
| AF1 | Alveolar fibroblast 1 (subtype of Matrix Fibroblast) |
| AF2 | Alveolar fibroblast 2 (subtype of Matrix Fibroblast) |
| ASMC | Airway Smooth Muscle (subtype of Mesenchymal cell) |
| Chondrocyte | Chondrocytes (subtype of Mesenchyme) |
| Mesothelial | Mesothelium (subtype of Mesenchyme) |
| Pericyte | Pericyte (subtype of Mesenchymal Cell) |
| SCMF | Secondary crest myofibroblast cell (subtype of Mesenchymal Cell) |
| VSMC | Vascular smooth muscle (subtype of Mesenchymal Cell) |
| **Immune Cells** | **Cell Type** |
| AM | Alveolar macrophage (subtype of Myeloid) |
| B | B Lymphocyte (subtype of Lymphocyte) |
| CD4_T | CD4+ T Cell (subtype of Immune) |
| CD8_T | CD8+ T Cell (subtype of Immune) |
| cDC1 | Classical DC subset 1 (subtype of Dendritic cell) |
| cDC2 | Classical DC subset 2 (subtype of Dendritic cell) |
| ILC | Innate lymphoid cells (subtype of Lymphocyte) |
| IM | Interstitial macrophage (subtype of Myeloid) |
| iMON | Inflammatory monocytes (subtype of Monocyte) |
| maDC | maDC, a highly specific marker of mature Dendritic cells (subtype of Myeloid) |
| Mast/Basophils | Mast (subtype of Myeloid) |
| Neutrophil | Neutrophil (subtype of Myeloid) |
| NK | Natural Killer Cell (subtype of Lymphocyte) |
| pDC | Plasmacytoid DC (subtype of Dendritic cell) |
| Plasma | Plasma Cell (subtype of Immune) |
| Platelet | Megakaryocyte/Platelet (subtype of Immune Cell) |
| pMON | Patrolling monocytes (subtype of Monocyte) |
| Treg | T Regulatory Cell (subtype of Immune) |

#### B. Color-coded human lung cell types

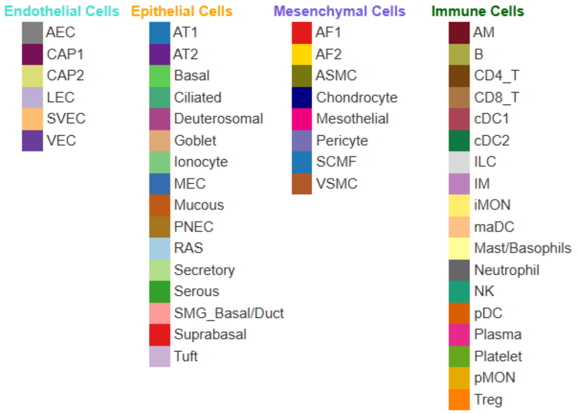

#### Supplemental Figure 4.

**A**

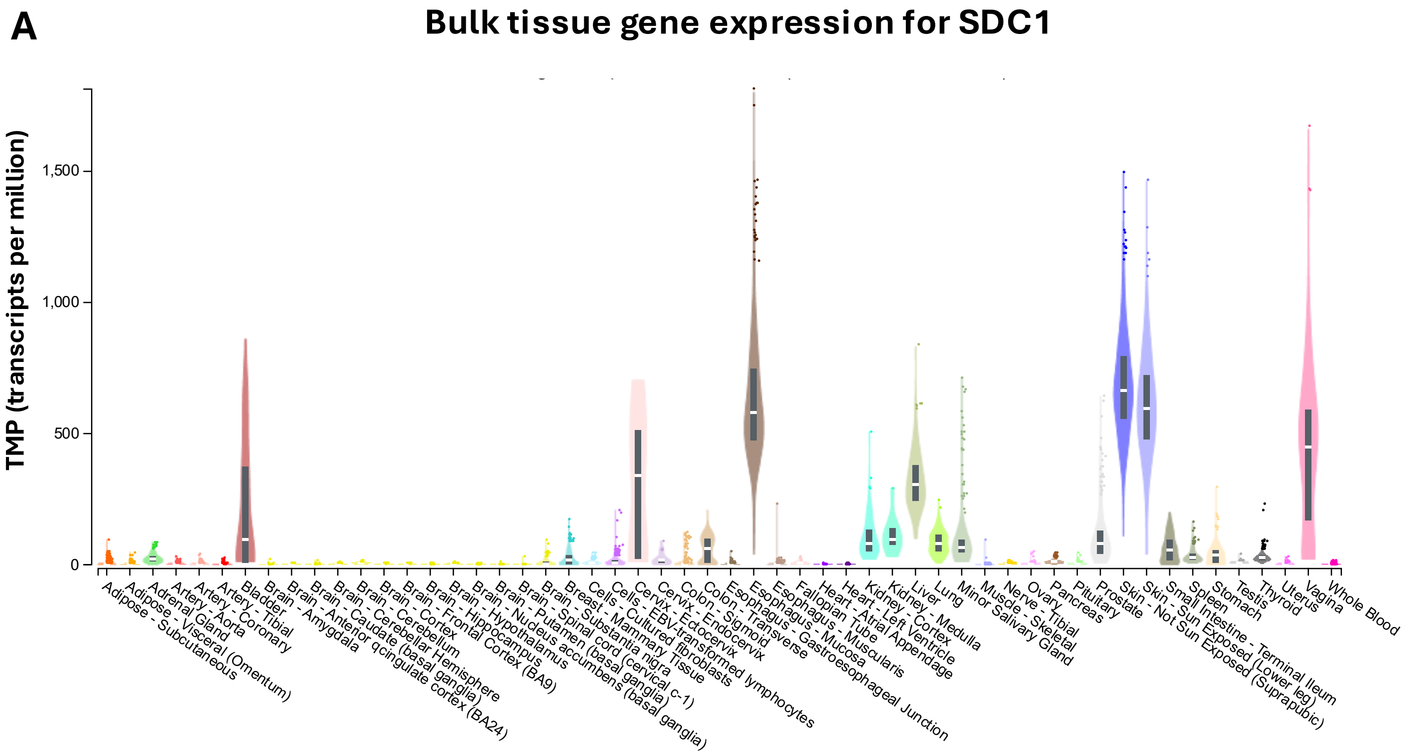


**B**

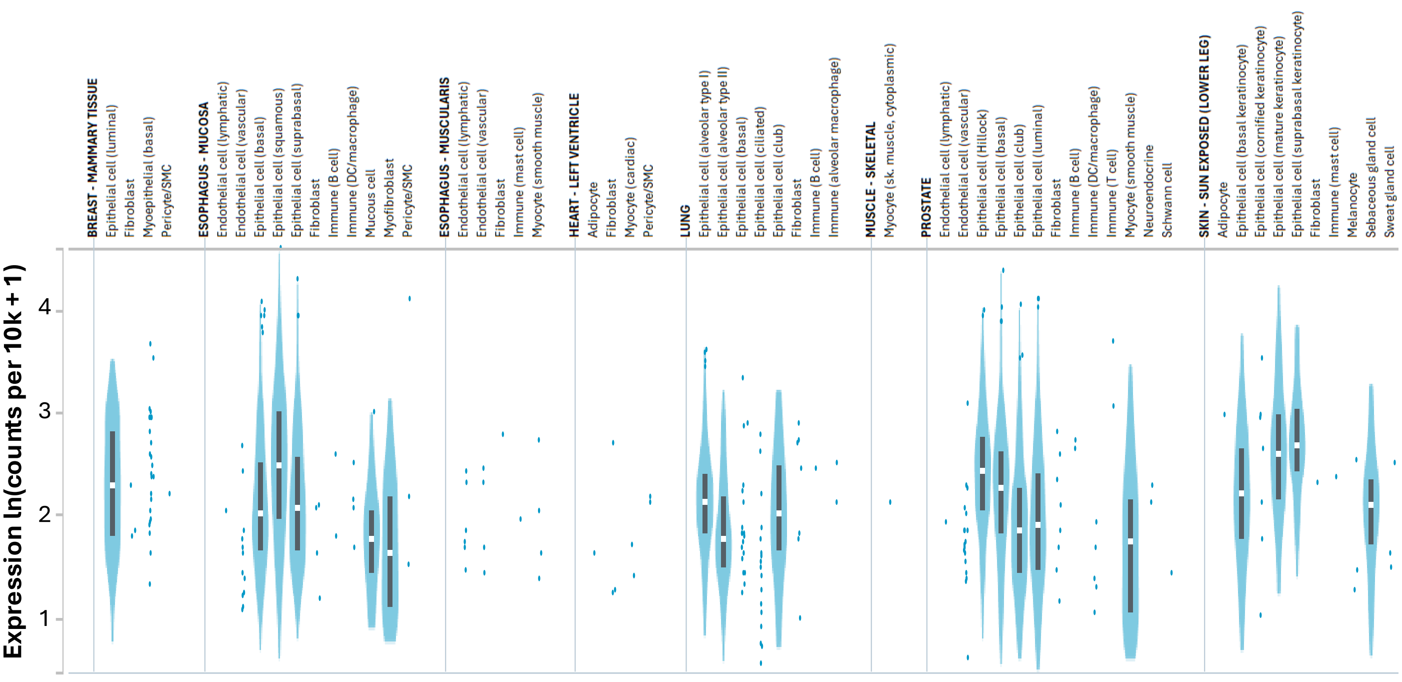

#### Supplemental Figure 5.

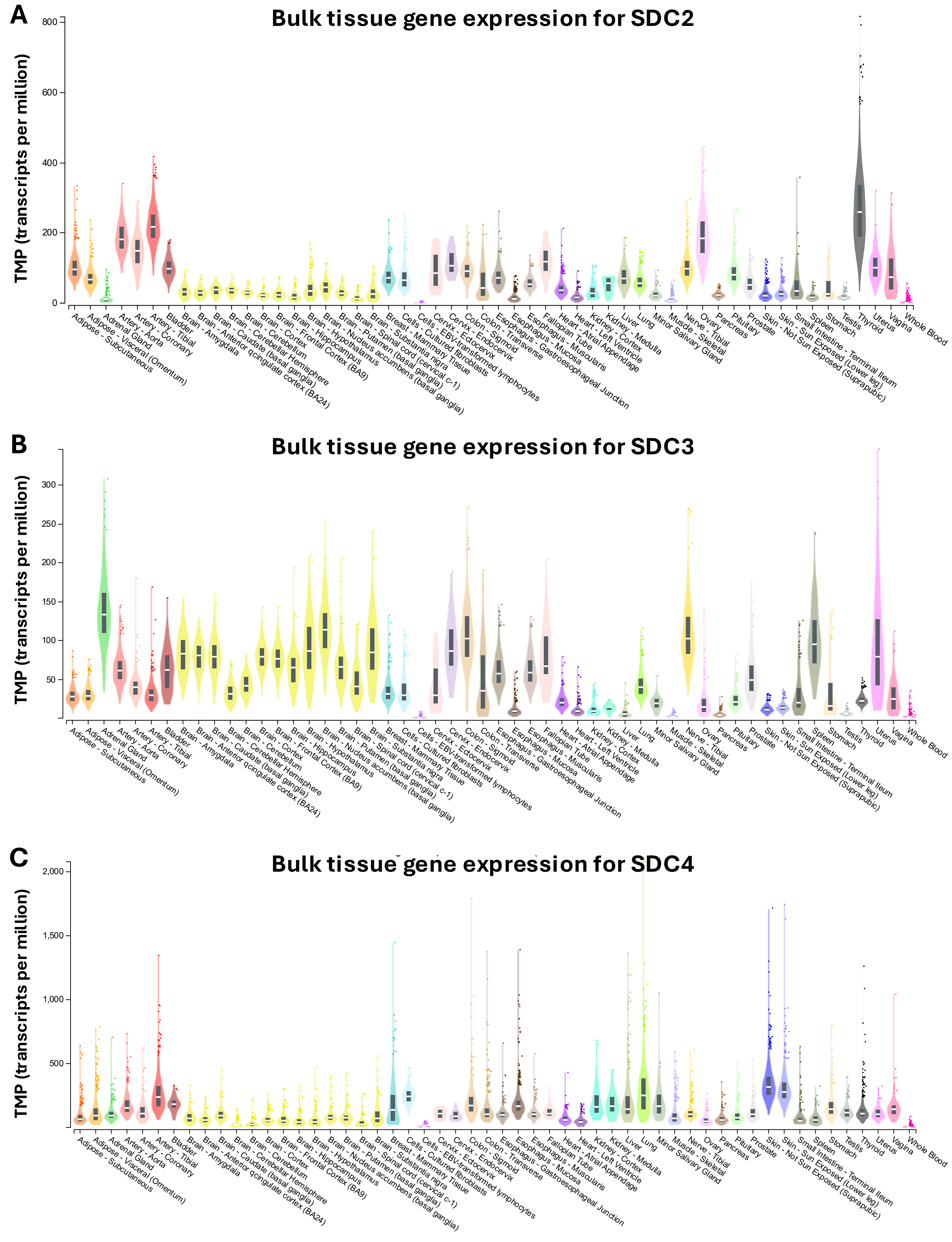

### Supplemental Figure 6.

**A**

#### Single-cell gene expression for SDC2

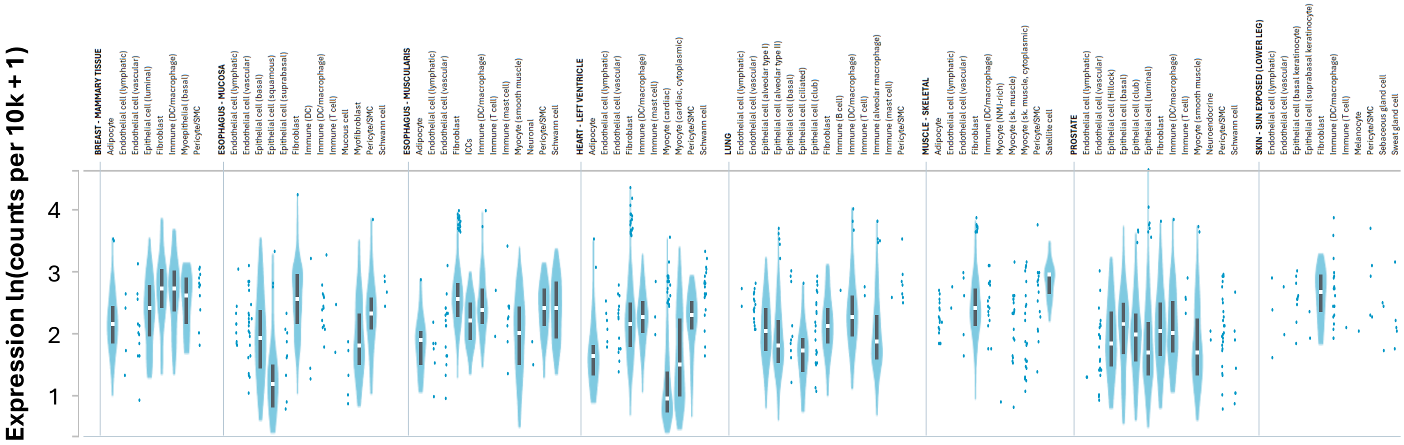


**B**

#### Single-cell gene expression for SDC3

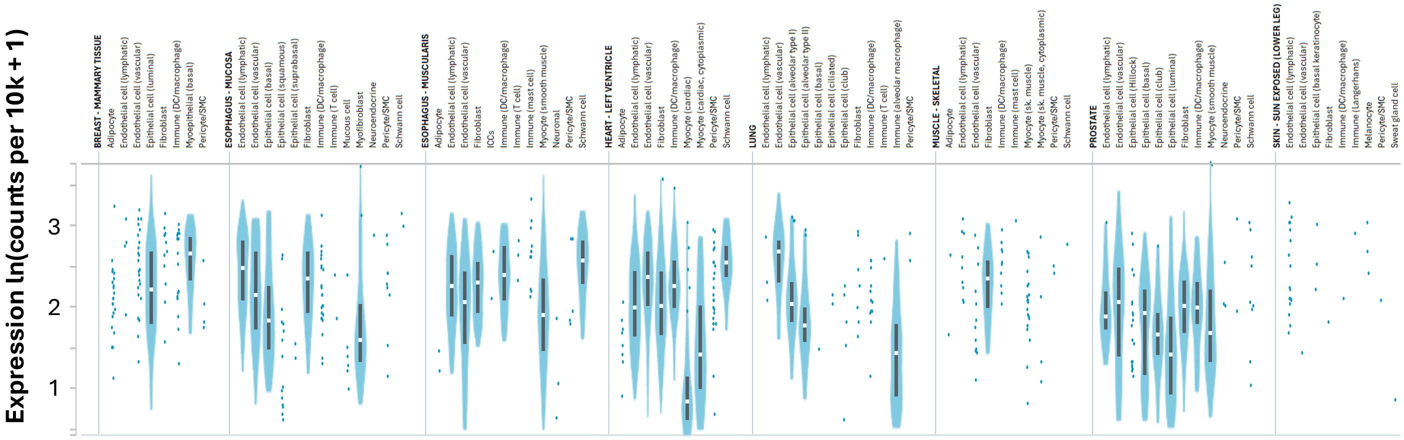


**C**

#### Single-cell gene expression for SDC4

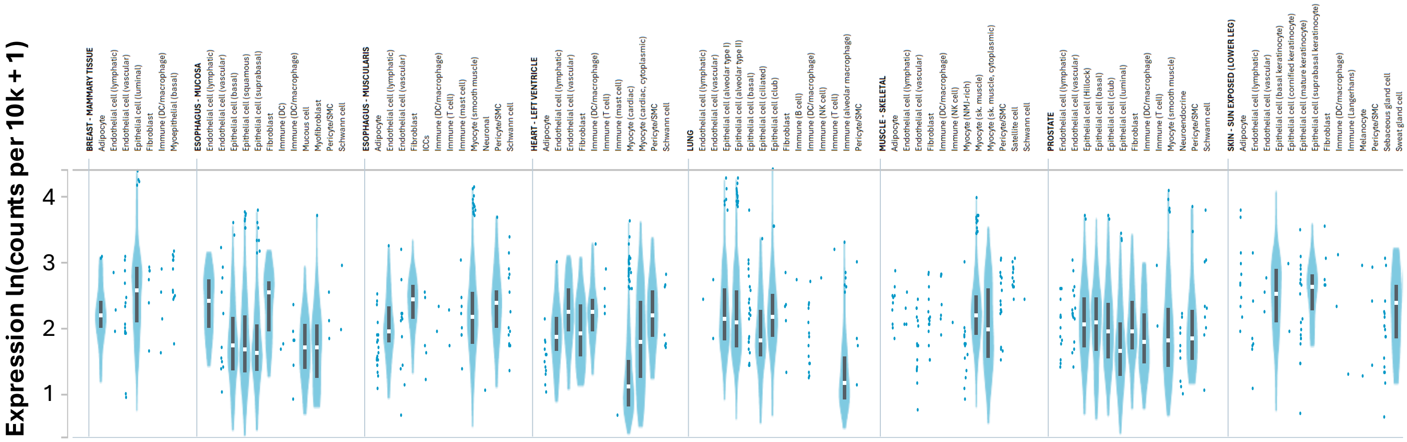

#### Supplemental Figure 7.

**Number of healthy donors: 175**
**Number of cells: 1,077,241**
**Number of tissues (organs): 20**

| | Tissue type | # cells | | Cell type | # cells |
|---|---|---|---|---|---|
| 1 | appendix | 2,199 | 1 | B and B plasma | 149,240 |
| 2 | ascending colon | 21,724 | 2 | Endothelial | 52,804 |
| 3 | buccal mucosa | 28,724 | 3 | Epithelial | 371,724 |
| 4 | caecum | 18,371 | 4 | Mesenchymal | 270,956 |
| 5 | colon | 156,256 | 5 | Myeloid | 33,213 |
| 6 | descending colon | 5,581 | 6 | Neural | 23,121 |
| 7 | duodenum | 144,244 | 7 | T and NK cells | 176,183 |
| 8 | gingival mucosa | 42,813 | | | |
| 9 | ileum | 214,737 | | | |
| 10 | intestine | 2,151 | | | |
| 11 | jejunum | 12,199 | | | |
| 12 | MLN | 35,312 | | | |
| 13 | oesophagus | 116,866 | | | |
| 14 | periodontium | 1,975 | | | |
| 15 | rectum | 18,950 | | | |
| 16 | salivary gland | 24,094 | | | |
| 17 | sigmoid colon | 28,503 | | | |
| 18 | small intestine | 29,473 | | | |
| 19 | stomach | 143,803 | | | |
| 20 | transverse colon | 29,266 | | | |
| | total # cells | 1,077,241 | | | 1,077,241 |

### Supplemental Figure 8.

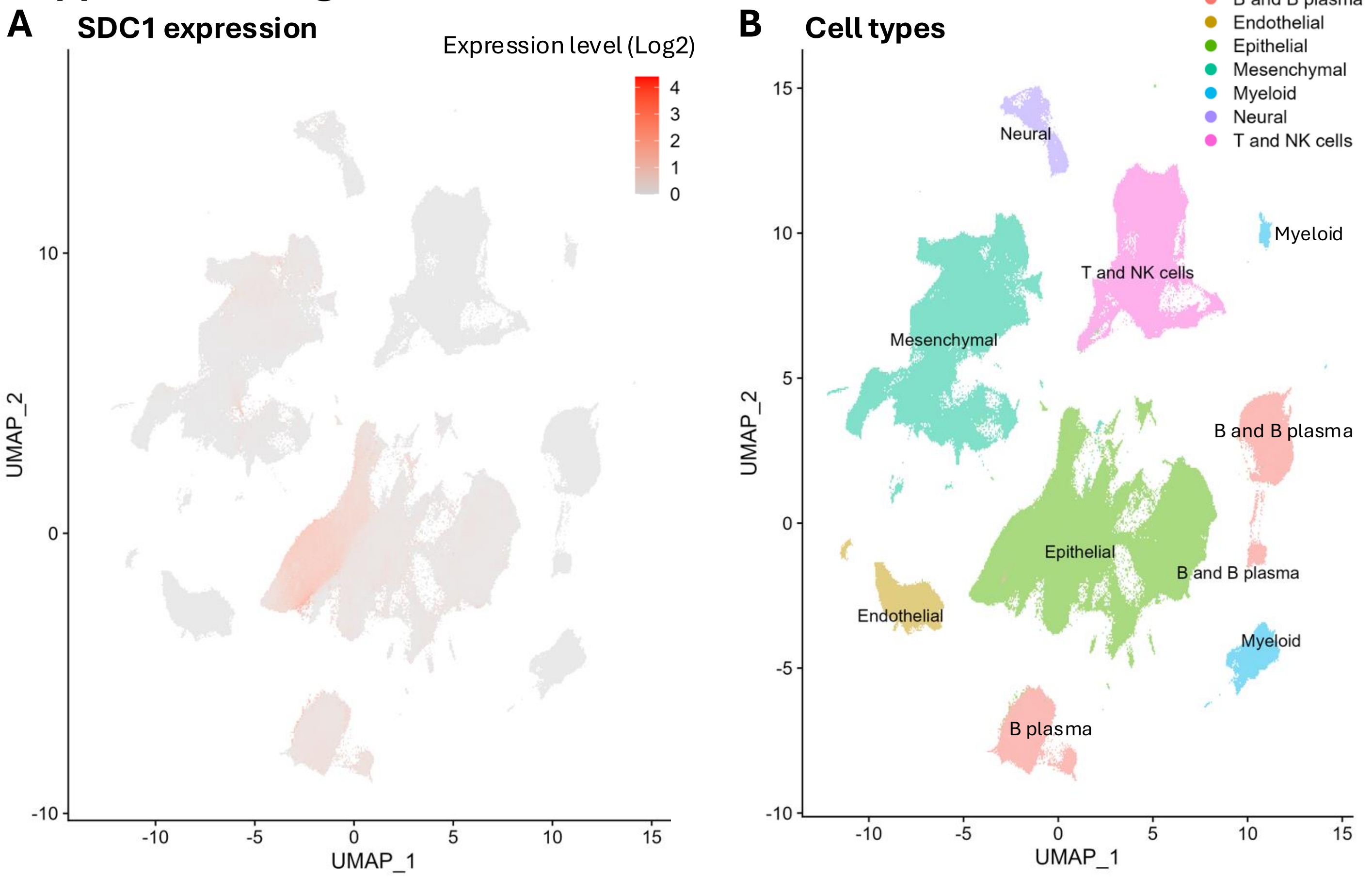


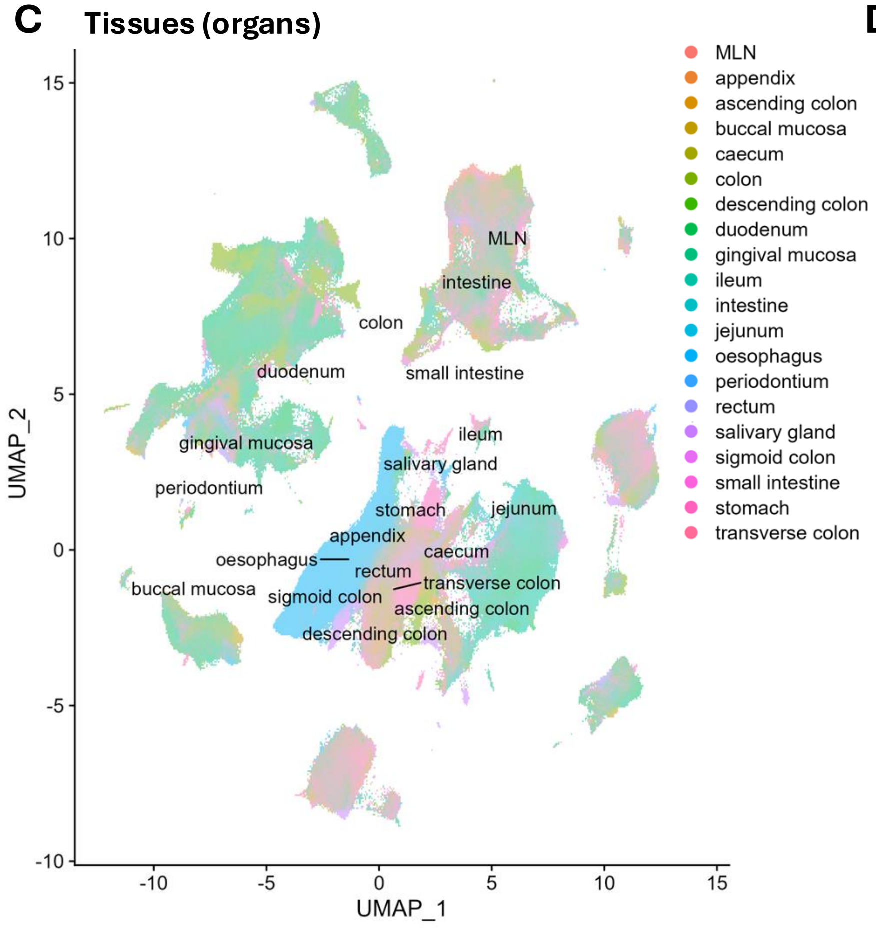


**D**

| Cell types | # Cells | % Cells (SDC1) | Mean Expr (SDC1) |
|---|---|---|---|
| **B plasma** | | | |
| B_plasma_IgA1 | 54,360 | 43.65 | 0.44 |
| B_plasma_IgA2 | 4,888 | 43.54 | 0.48 |
| B_plasma_IgG | 7,033 | 43.00 | 0.46 |
| B_plasma_IgM | 4,922 | 35.45 | 0.35 |
| B_plasmablast | 465 | 37.20 | 0.36 |
| **Endothelial** | | | |
| EC_arterial_1 | 4,780 | 0.19 | 0.00 |
| EC_arterial_2 | 3,507 | 0.14 | 0.00 |
| EC_capillary | 11,035 | 0.43 | 0.00 |
| EC_cycling | 1,070 | 1.50 | 0.01 |
| EC_lymphatic | 4,248 | 1.04 | 0.01 |
| EC_venous | 28,164 | 0.75 | 0.01 |
| **Epithelial** | | | |
| Basal | 13,722 | 46.71 | 0.58 |
| Basal_cycling | 2,252 | 69.32 | 0.76 |
| Epithelial_cycling_G2M | 15,948 | 57.11 | 0.60 |
| Epithelial_cycling_S | 15,166 | 53.92 | 0.49 |
| Epithelial_progenitor: | | | |
| Distal_progenitor | 2,765 | 27.31 | 0.16 |
| Distal_progenitor_PRAC1 | 323 | 56.04 | 0.29 |
| Epithelial_stem | 10,520 | 25.38 | 0.16 |
| Deep crypt secretory: | | | |
| DCS_MUC17 | 231 | 48.48 | 0.43 |
| DCS_MUC17_cycling | 107 | 41.12 | 0.27 |
| Gland_basal | 735 | 46.80 | 0.59 |
| Gland_duct | 254 | 38.19 | 0.38 |
| Gland_fetal | 248 | 92.34 | 1.31 |
| Gland_mucous | 53 | 15.09 | 0.15 |
| Goblet | 17,475 | 23.48 | 0.20 |
| Goblet_cycling | 1,494 | 43.64 | 0.28 |
| Goblet_progenitor | 1,935 | 18.76 | 0.20 |
| Keratinocyte: | | | |
| fetal | 125 | 95.20 | 1.38 |
| inflammatory | 284 | 60.56 | 0.50 |
| outer | 14,615 | 81.09 | 1.09 |
| stratified | 63,400 | 89.56 | 1.38 |

### Supplemental Figure 9.

**A** SDC2 expression

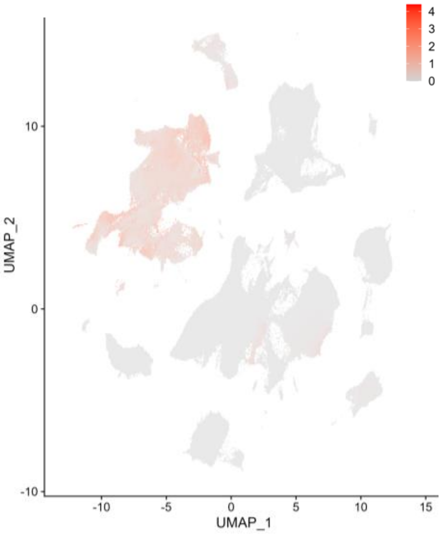


**B** SDC3 expression

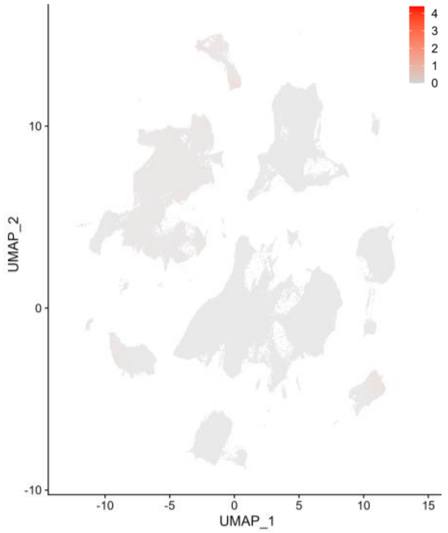


**C** SDC4 expression

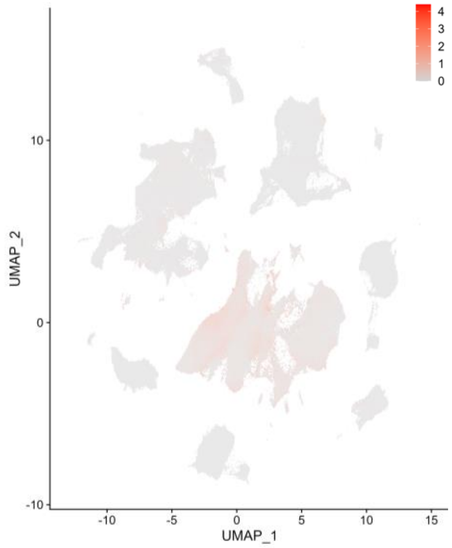


**D**

| Cells types | # Cells | % Cells (SDC2) | Mean Expr (SDC2) |
|---|---|---|---|
| **B plasma** | | | |
| B_plasma_IgA1 | 54,360 | 0.11 | 0.00 |
| B_plasma_IgA2 | 4,888 | 0.04 | 0.00 |
| B_plasma_IgG | 7,033 | 0.24 | 0.00 |
| B_plasma_IgM | 4,922 | 0.04 | 0.00 |
| B_plasmablast | 465 | 0.00 | 0.00 |
| **Endothelial** | | | |
| EC_arterial_1 | 4,780 | 0.86 | 0.01 |
| EC_arterial_2 | 3,507 | 0.68 | 0.01 |
| EC_capillary | 11,035 | 1.24 | 0.01 |
| EC_cycling | 1,070 | 4.67 | 0.06 |
| EC_lymphatic | 4,248 | 1.74 | 0.02 |
| EC_venous | 28,164 | 1.08 | 0.01 |
| **Epithelial** | | | |
| Keratinocyte | | | |
| fetal | 125 | 0.00 | 0.00 |
| inflammatory | 284 | 1.06 | 0.01 |
| outer | 14,615 | 0.09 | 0.00 |
| stratified | 63,400 | 0.25 | 0.00 |
| **Mesenchymal** | | | |
| Fibroblast | | | |
| Crypt_fibroblast_PI16 | 41,313 | 49.53 | 0.80 |
| Cycling_fibroblast | 18,017 | 67.43 | 1.00 |
| Lamina_propria_fibroblast | 37,736 | 52.20 | 0.85 |
| Oesophagus_fibroblast | 682 | 26.69 | 0.46 |
| Oral_mucosa_fibroblast | 4,378 | 57.10 | 0.88 |
| Rectum_fibroblast | 1,284 | 64.88 | 1.11 |
| Villus_fibroblast_F3 | 20,819 | 73.87 | 1.37 |
| Intestinal_Cajal_cell (ICC) | 923 | 24.16 | 0.27 |
| Lymphoid_stromal_cell | | | |
| Fibroblast_reticular | 2,562 | 44.03 | 0.63 |
| Follicular_DC | 135 | 15.56 | 0.11 |
| Mesenchymal_LTO | 353 | 49.01 | 0.57 |
| Mesoderm_1 | 30022 | 49.27 | 0.62 |
| Mesoderm_2 | 53169 | 46.80 | 0.71 |
| Mesothelium | 2784 | 58.05 | 0.57 |
| Myoblast/myocyte | 133 | 65.41 | 0.84 |
| Myofibroblast | 15509 | 52.91 | 0.84 |
| Pericyte | | | |
| Angiogenic_pericyte | 849 | 89.16 | 1.80 |
| Immature_pericyte | 2,716 | 59.43 | 1.07 |
| Immune_recruiting_pericyte | 363 | 60.61 | 1.04 |
| Pericyte | 6,974 | 62.42 | 1.12 |
| Smooth_muscle | | | |
| SMC_CAPN3 | 6,293 | 40.93 | 0.48 |
| SMC_CAPN3_cycling | 442 | 61.31 | 0.65 |
| SMC_PPLP2 | 14,522 | 60.09 | 0.70 |
| Vascular_smooth_muscle | 8,978 | 41.58 | 0.68 |
| **Neural** | | | |
| Neuron | | | |
| Branch_A_inhibtory_motor_neuron | 861 | 25.55 | 0.33 |
| Branch_A_interneuron | 684 | 19.44 | 0.18 |
| Branch_A_primary_afferent_neuron | 1,013 | 56.37 | 0.66 |
| Branch_B_excitatory_motor_neuron | 2,955 | 13.91 | 0.17 |
| Branch_B_primary_afferent_neuron | 1,158 | 29.02 | 0.29 |
| Neuron_progenitor | | | |
| Enteric_neural_crest_cycling | 4,107 | 27.05 | 0.24 |
| Glial/Enteric_neural_crest | 5,024 | 16.12 | 0.21 |
| Glial_1 | 1,069 | 14.03 | 0.15 |
| Glial_2 | 1,501 | 29.98 | 0.42 |
| Glial_3 | 2,036 | 23.92 | 0.28 |
| Neuroblast | 2,713 | 13.31 | 0.15 |

**E**

| Cells types | # Cells | % Cells (SDC3) | Mean Expr (SDC3) |
|---|---|---|---|
| **B plasma** | | | |
| B_plasma_IgA1 | 54,360 | 0.06 | 0.00 |
| B_plasma_IgA2 | 4,888 | 0.10 | 0.00 |
| B_plasma_IgG | 7,033 | 0.14 | 0.00 |
| B_plasma_IgM | 4,922 | 0.02 | 0.00 |
| B_plasmablast | 465 | 0.22 | 0.00 |
| **Endothelial** | | | |
| EC_arterial_1 | 4,780 | 14.98 | 0.17 |
| EC_arterial_2 | 3,507 | 11.61 | 0.13 |
| EC_capillary | 11,035 | 9.74 | 0.09 |
| EC_cycling | 1,070 | 11.50 | 0.07 |
| EC_lymphatic | 4,248 | 20.69 | 0.24 |
| EC_venous | 28,164 | 3.10 | 0.03 |
| **Epithelial** | | | |
| Basal | 2,252 | 2.09 | 0.01 |
| Basal_cycling | 13,722 | 1.39 | 0.01 |
| Epithelial_cycling_G2M | 15,948 | 4.46 | 0.02 |
| Epithelial_cycling_S | 15,166 | 5.83 | 0.02 |
| Epithelial_progenitor: | | | |
| Distal_progenitor | 2,765 | 11.25 | 0.06 |
| Distal_progenitor_PRAC1 | 323 | 14.55 | 0.05 |
| Epithelial_stem | 10,520 | 2.95 | 0.01 |
| Deep crypt secretory: | | | |
| DCS_MUC17 | 231 | 5.63 | 0.03 |
| DCS_MUC17_cycling | 107 | 13.08 | 0.06 |
| Gland_basal | 735 | 0.54 | 0.00 |
| Gland_duct | 254 | 4.33 | 0.03 |
| Gland_fetal | 248 | 0.00 | 0.00 |
| Gland_mucous | 53 | 0.00 | 0.00 |
| Keratinocyte: | | | |
| fetal | 125 | 2.40 | 0.00 |
| inflammatory | 284 | 1.06 | 0.01 |
| outer | 14,615 | 0.01 | 0.00 |
| stratified | 63,400 | 0.10 | 0.00 |
| **Neural** | | | |
| Neuron | | | |
| Branch_A_inhibtory_motor_neuron | 861 | 26.48 | 0.26 |
| Branch_A_interneuron | 684 | 36.84 | 0.37 |
| Branch_A_primary_afferent_neuron | 1,013 | 47.58 | 0.49 |
| Branch_B_excitatory_motor_neuron | 2,955 | 15.19 | 0.16 |
| Branch_B_primary_afferent_neuron | 1,158 | 50.17 | 0.54 |
| Neuron_progenitor | | | |
| Enteric_neural_crest_cycling | 4107 | 17.09 | 0.14 |
| Glial/Enteric_neural_crest | 5024 | 11.92 | 0.16 |
| Glial_1 | 1,069 | 23.39 | 0.27 |
| Glial_2 | 1,501 | 28.65 | 0.38 |
| Glial_3 | 2,036 | 12.48 | 0.13 |
| Neuroblast | 2,713 | 8.00 | 0.09 |

**F**

| Cell types | # Cells | % Cells (SDC4) | Mean Expr (SDC4) |
|---|---|---|---|
| **B plasma** | | | |
| B_plasma_IgA1 | 54,360 | 0.61 | 0.00 |
| B_plasma_IgA2 | 4,888 | 0.55 | 0.00 |
| B_plasma_IgG | 7,033 | 0.97 | 0.01 |
| B_plasma_IgM | 4,922 | 0.65 | 0.01 |
| B_plasmablast | 465 | 0.86 | 0.01 |
| **Endothelial** | | | |
| EC_arterial_1 | 4,780 | 1.28 | 0.02 |
| EC_arterial_2 | 3,507 | 0.97 | 0.01 |
| EC_capillary | 11,035 | 0.64 | 0.01 |
| EC_cycling | 1,070 | 2.06 | 0.02 |
| EC_lymphatic | 4,248 | 12.66 | 0.15 |
| EC_venous | 28,164 | 0.79 | 0.01 |
| **Epithelial** | | | |
| Basal | 2,252 | 39.74 | 0.39 |
| Basal_cycling | 13,722 | 30.07 | 0.36 |
| Epithelial_cycling_G2M | 15,948 | 48.94 | 0.42 |
| Epithelial_cycling_S | 15,166 | 50.07 | 0.39 |
| Epithelial_progenitor: | | | |
| Distal_progenitor | 2,765 | 53.60 | 0.49 |
| Distal_progenitor_PRAC1 | 323 | 64.40 | 0.45 |
| Epithelial_stem | 10,520 | 40.95 | 0.31 |
| Deep crypt secretory: | | | |
| DCS_MUC17 | 231 | 55.41 | 0.51 |
| DCS_MUC17_cycling | 107 | 36.45 | 0.23 |
| Gland_basal | 735 | 41.36 | 0.56 |
| Gland_duct | 254 | 68.90 | 0.91 |
| Gland_fetal | 248 | 85.08 | 0.99 |
| Gland_mucous | 53 | 35.85 | 0.36 |
| Goblet | 17,475 | 32.69 | 0.34 |
| Goblet_cycling | 1,494 | 41.50 | 0.27 |
| Goblet_progenitor | 1,935 | 25.94 | 0.28 |
| Keratinocyte: | | | |
| fetal | 125 | 93.60 | 1.16 |
| inflammatory | 284 | 39.08 | 0.32 |
| outer | 14,615 | 39.75 | 0.30 |
| stratified | 63,400 | 61.57 | 0.62 |

#### Supplemental Figure 10.

Number of healthy donors: 4
Number of cells: All neurons 2,480,956;
All non-neuronal cells: 888,263
Number of tissues: 10

**A**

all neuron

| | Locations | # cells | | Cell type | # cells |
|---|---|---|---|---|---|
| 1 | cerebellum | 102,944 | 1 | Neuron | 2,477,203 |
| 2 | cerebral cortex | 1,156,201 | 2 | Leukocyte | 3,753 |
| 3 | cerebral nuclei | 372,383 | | | |
| 4 | hippocampal formation | 237,967 | | | |
| 5 | hypothalamus | 110,280 | | | |
| 6 | midbrain | 127,849 | | | |
| 7 | myelencephalon | 47,352 | | | |
| 8 | pons | 107,041 | | | |
| 9 | spinal cord | 6,038 | | | |
| 10 | thalamic complex | 212,901 | | | |
| | total # cells | 2,480,956 | | | 2,480,956 |

**B**

all non-neuron

| | Locations | # cells | | Cell type | # cells |
|---|---|---|---|---|---|
| 1 | cerebellum | 49,003 | 1 | Bergmann glial cell | 8,041 |
| 2 | cerebral cortex | 148,874 | 2 | astrocyte | 155,025 |
| 3 | cerebral nuclei | 150,926 | 3 | central nervous system macrophage | 91,838 |
| 4 | hippocampal formation | 109,980 | 4 | choroid plexus epithelial cell | 7,689 |
| 5 | hypothalamus | 24,191 | 5 | endothelial cell | 5,165 |
| 6 | midbrain | 121,456 | 6 | ependymal cell | 5,882 |
| 7 | myelencephalon | 38,135 | 7 | fibroblast | 9,156 |
| 8 | pons | 97,127 | 8 | oligodendrocyte | 494,966 |
| 9 | spinal cord | 18,152 | 9 | oligodendrocyte precursor cell | 105,734 |
| 10 | thalamic complex | 130,419 | 10 | pericyte | 3,693 |
| | | | 11 | vascular associated smooth muscle cell | 1,074 |
| | total # cells | 888,263 | | | 888,263 |

#### Supplemental Figure 11.

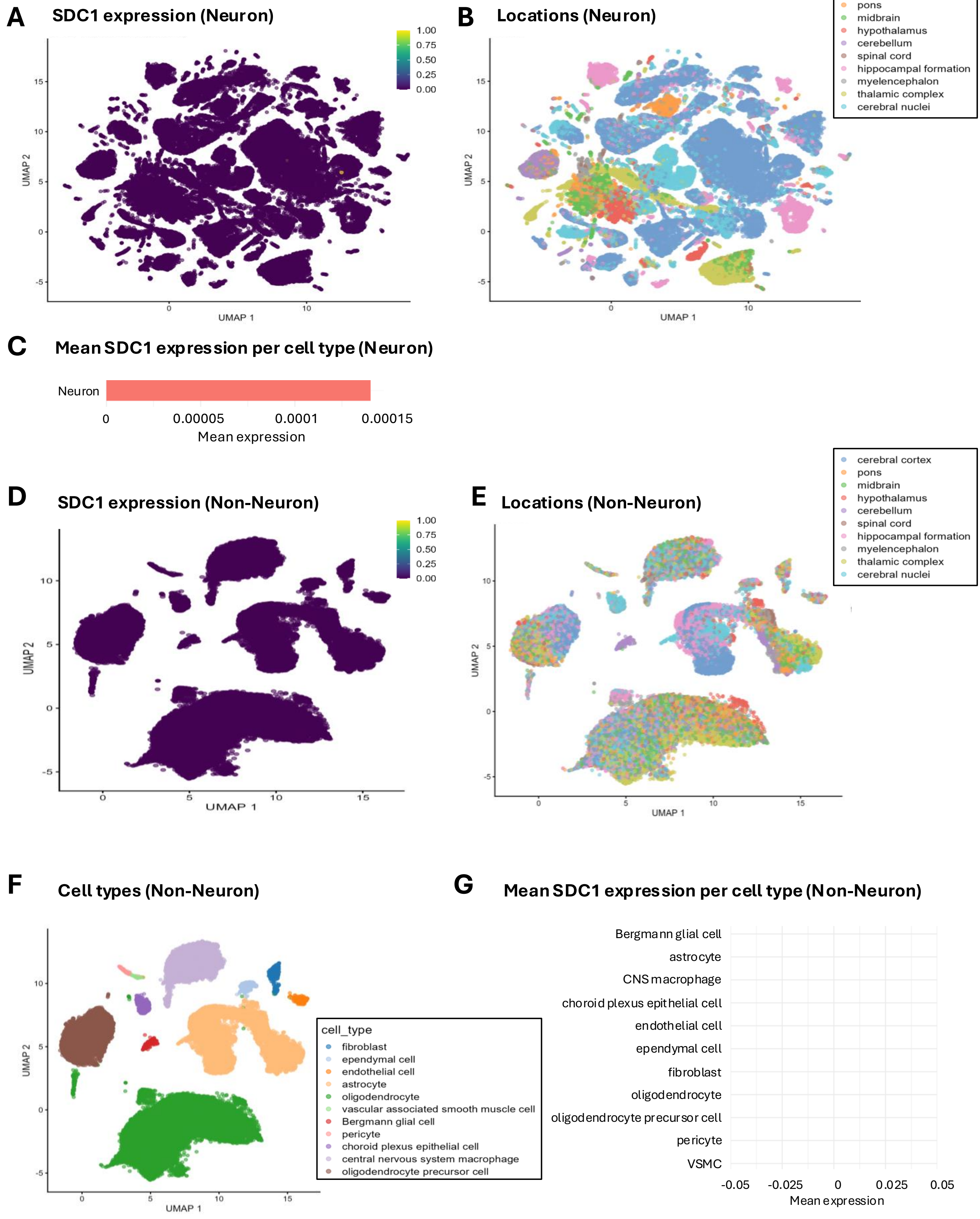

### Supplemental Figure 12.

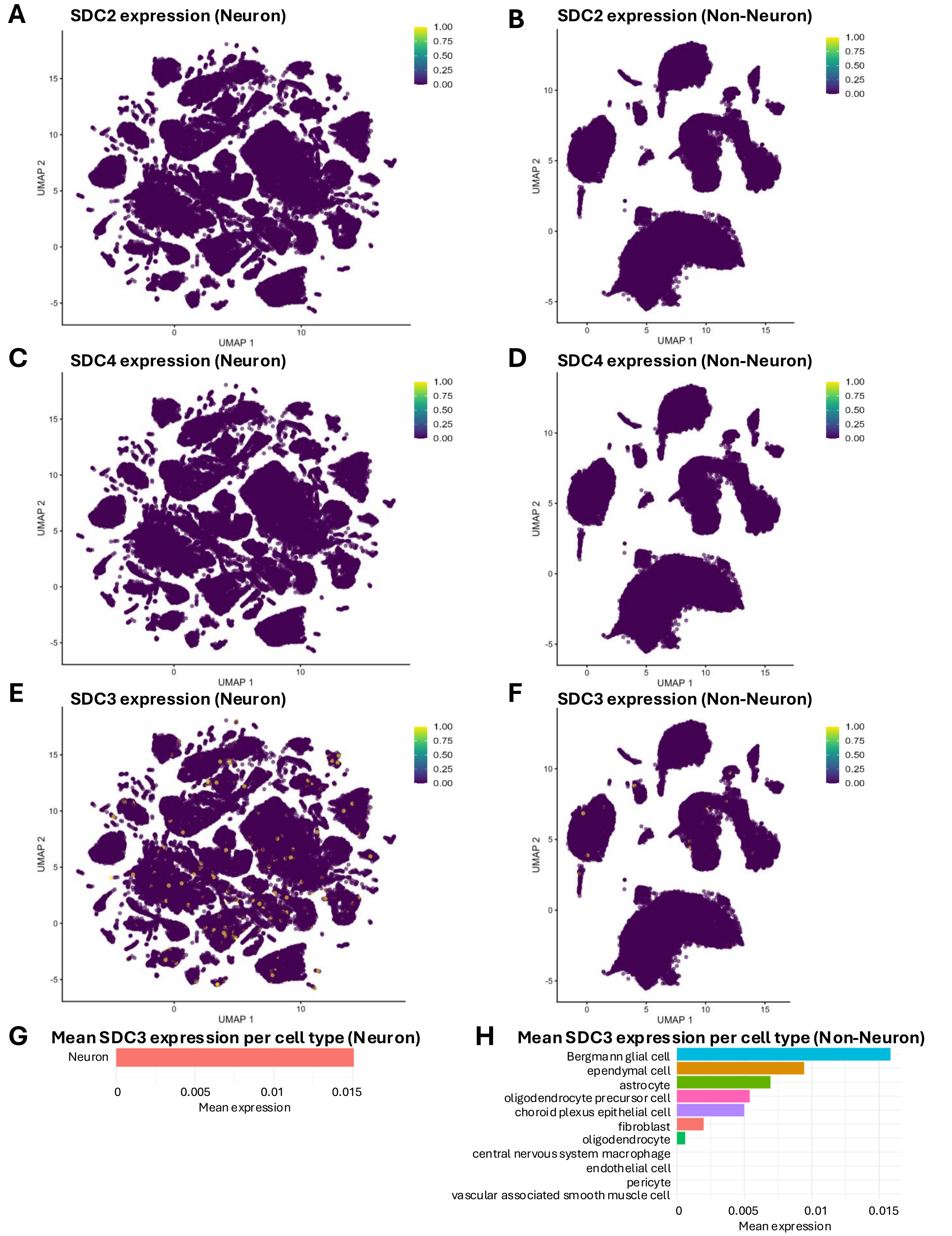
